# Adiponectin Receptor Fragmentation in Mouse Models of Type 1 and Type 2 Diabetes

**DOI:** 10.1101/849000

**Authors:** Dylan Frabutt, Natalie Stull, Annie R. Pineros, Sarah A. Tersey, Donalyn Scheuner, Teresa L. Mastracci, Michael J. Pugia

**Affiliations:** Indiana Biosciences Research Institute, Indianapolis IN; Center for Diabetes and Metabolic Diseases, Indiana University School of Medicine, Indianapolis IN, USA

## Abstract

The protein hormone adiponectin regulates glucose and fatty acid metabolism by binding to two PAQR-family receptors (AdipoR1 and AdipoR2). Both receptors feature a C-terminal segment which is released by proteolysis to form a freely-circulating C-terminal fragment (CTF) found in the plasma of normal individuals but not in some undefined diabetes patients. The AdipoR1-CTF_344-376_ is a competitive inhibitor of tumor necrosis factor α cleavage enzyme (TACE) but it contains a shorter peptide domain (AdipoR1 CTF_351-362_) that is a strong non-competitive inhibitor of insulin-degrading enzyme (IDE). The link between adiponectin receptor fragmentation and diabetes pathology is unclear but could lead to new therapeutic strategies. We therefore investigated physiological variations in the concentrations of CTF in non-obese diabetic (NOD/ShiLtJ) mice and C57BL/6 mice with diet-induced obesity (DIO) as models of diabetes types 1 and 2, respectively. We tested for changes in adiponectin receptor signaling, immune responses, disease progression, and the abundance of neutralizing autoantibodies. Finally, we administered exogenous AdipoR1-CTF peptides either containing or lacking the IDE-binding domain. We observed the more pronounced CTF shedding in the TACE-active NOD mice, which represents an inflammatory autoimmune phenotype, but fragmentation was also observed to a lesser extent in the DIO model. Autoantibodies to CTF were detected in both models. Neither exogenous CTF peptide affected IgG-CTF plasma levels, body weight or the conversion of NOD mice to diabetes. The pattern of AdipoR1 fragmentation and autoantibody production under physiological conditions of aging, DIO, and autoimmune diabetes therefore provides insight into the association adiponectin biology and diabetes.

## Introduction

An association between inflammation, obesity and adiponectin biology is emerging in studies of Type 1 diabetes (T1D) and its complications, promoting the investigation of adiponectin receptor biology in models of both T1D and Type 2 diabetes (T2D). Total and high-molecular-weight adiponectin levels are elevated in the serum of T1D patients, but there are lower levels of adiponectin receptor expression on monocytes and antigen-presenting cells, which may be responsible for increased inflammation.^1-5^ In contrast, lower serum levels of adiponectin are associated with T2D, cardiovascular disease, and insulin resistance, although the levels increase when T2D patients undergo post-gastric bypass surgery for visceral fat reduction.^6,7^ The downregulation of adiponectin receptor expression in T1D patients is associated with the proliferation of primary CD4^+^ T-cells in the presence of islet lysate independent of adiponectin treatment, whereas healthy control T-cells are inhibited by adiponectin.^8^

Both isoforms of the adiponectin receptor (AdipoR1 and AdipoR2) contain an extracellular, C-terminal sequence. This is released by proteolytic degradation to yield a C-terminal fragment (CTF) that can be detected in human, murine and rat peripheral blood.^9^ The role of this peptide is unclear, but it features an insulin-degrading enzyme (IDE)-binding domain.^9^ Complexes of autoantibodies interacting with CTF have been discovered in humans and rodents.^9^ In a preliminary study comparing murine models of T1D and T2D, we observed differences in the levels of CTF release. Specifically, higher levels of CTF were associated with pancreatic tissue with higher levels of tumor necrosis factor α cleavage enzyme (TACE, also known as ADAM-17/MMP-17) activity (**Supplemental Report 1**).

## Experimental goal

We propose that a better understanding of adiponectin receptor fragmentation will provide insight into T1D pathology and could help to identify new therapeutic strategies. We therefore investigated physiological variations in the concentrations of CTF in non-obese diabetic (NOD/ShiLtJ) mice and C57BL/6 mice with diet-induced obesity (DIO). We also tested for changes in adiponectin receptor signaling, immune responses, disease progression, and the abundance of neutralizing autoantibodies. Finally, we administered exogenous AdipoR1-CTF peptides either containing or lacking the IDE-binding domain to further assess the function of this circulating peptide fragment.

## Experimental section

### Materials

A mouse monoclonal antibody (mAb) specific for AdipoR1-CTF_351-375_ (clone 444-1D12-1H7) was produced in hybridoma cell lines (ATCC) and was purified using a protein G column (IBRI, Indianapolis, IN). A rabbit mAb specific for AdipoR1-CTF_351-362_ (Epitomics clone SAT 5.2, 0.5 mg/mL) was obtained as purified material (SISCAPA Technologies). A rabbit polyclonal antibody (pAb) specific for AdipoR1-CTF_351-375_ was covalently bound to alkaline phosphatase (ALP; Siemens Healthcare) and was used in sandwich assays for free CTF.^9^ The synthetic peptides for inhibitory AdipoR1-CTF_351-375_ (25-mer) and non-inhibitory AdipoR1-CTF_358-375_ (18-mer) were acquired from Celtek Biosciences. The non-competitive inhibition of IDE was confirmed for AdipoR1-CTF_351-375_ but not AdipoR1-CTF_358-375_ as previously reported,^1^ and ∼90% of IDE activity is inhibited at a CTF concentration of 10 µg/mL (**Supplemental Report 2**). AdipoR1-CTF mAb clone 444 was conjugated directly to FITC (0.5 mg/mL) for FACS-based immunocytochemical (ICC) and immunohistochemical (IHC) analysis. We also obtained the following antibodies from the suppliers as shown: AMPKα rabbit (AbCam, 2532S), P-AMPKα (T172) rabbit (AbCam, 2535S), Akt rabbit (AbCam 9272S), P-Akt (S473) mouse mAb (AbCam, 4051S), β-actin mouse mAb (Thermo Fischer Scientific, MA5-15739) and TACE/ADAM17 pAb (R&D Systems, AF9301).

## Methods

### Animal treatments

The Institutional Animal Care and Use Committee (IACUC) of the Indiana University School of Medicine, Indianapolis, IN, approved all animal procedures. Male C57BL/6J mice were used as a model of obesity-induced diabetes and showed a high incidence of diabetes after 12 weeks on a high-fat diet consisting of 60% kcal from fat.^10^ Male C57BL/6J mice maintained on a normal diet were used as the normal control group. Female NOD/ShiLtJ mice were maintained on a normal diet and were used as a model of non-obese diabetes. This strain is a well-characterized polygenic model of T1D featuring insulitis, leukocyte infiltration of the pancreatic islets, and pancreatic TACE expression.^11,12^ Female NOD/ShiLtJ mice show a higher incidence of diabetes by 16 weeks (∼80% in our facility). Male NOD/ShiLtJ mice were not used because they show a low incidence of diabetes (<20%) and lack pancreatic TACE expression.^12^ The strains are abbreviated to C57 and NOD hereafter.

Female C57 and NOD mice arrived at 4 weeks of age, and both strains began to present diabetes at 12 weeks. Blood glucose and body weight were measured weekly. After 13 days, the animals were sorted into groups based on plasma glucose levels and body weight, and sorting continued every week until day 210. After 14 and 105 days, or following termination due to diabetes, blood and tissue were collected from female C57 and NOD mice (n=5 for each subgroup) by cardiac puncture, saline perfusion and dissection of the pancreas, spleen, liver, white adipose, liver, brown adipose, brain and quad muscle tissues. The samples were fixed, embedded in OCT and sectioned, and slides were processed for ICC/IHC staining. Pancreas tissue was harvested from additional groups of n=5 mice for insulitis staining, as well as western blot analysis (whole pancreas, pancreas islets and spleens). Pancreatic lymph nodes and spleens were harvested from the same mice for FACS analysis of T-cell counts.^13,14^

C57 and NOD mice were treated weekly either with vehicle, AdipoR1-CTF_351-375_ or AdipoR1-CTF_358-375_ by intraperitoneal (IP) injections starting at 14 days (6 weeks of age) and continuing for up to 105 days (14 weeks of age). Compounds were administered at a dose of ∼1 mg/kg (125 µL of 200 µg/mL sterile saline per 25 g body weight) after dilution from 4 mg/mL DMF stocks (stored at 4°C) to 5% DMF in saline vehicle. In the first cohort, NOD female mice on the normal diet and C57 male mice on the high-fat diet were treated with vehicle, AdipoR1-CTF_351-375_, or TACE inhibitor, a glucose tolerance test (GTT) was performed at 6 and 14 weeks of age, and the animals were euthanized at 6 and 16 weeks of age (**Supplemental Report 1**). In the second cohort, NOD female mice on the normal diet were treated with vehicle, AdipoR1-CTF_351-375_ or AdipoR1-CTF_358-_ 375, whereas C57 female mice on the normal and high-fat diets were treated with vehicle alone. A GTT was performed at 14 weeks, and an intraperitoneal insulin tolerance test (IPITT) at 16 weeks. Animals were euthanized at 6, 16 and 24 weeks of age as was any mouse with a blood glucose reading >250 mg/dL for more than 2 consecutive days (Protocols flow charts are compared in **Supplemental Report 1**).

### GTT method

Mice were fasted for either 5 or 16 h (with full access to water) before IP injection with 1–2 g/kg glucose. We then took 1–2 µL blood samples by tail biopsy at 0, 10, 20, 30, 60, 90 and 120 min after glucose dosing. At three time points (0, 60 and 120 min), 40 µL of blood was collected for insulin and CTF measurements (20 µL per metabolite) using a single tail biopsy in each case (less than needed to obtain DNA for genotyping). Bleeding from the tail between sampling times was minimal due to clot formation. To obtain each sample, the clot was removed by wiping with an alcohol pad to remove the clot, and a drop of blood was encouraged to form by squeezing the tail from body to tip. The blood drop was picked up on a glucometer strip for analysis using an AlphaTrak 2 blood glucose monitor.

### IPITT method

Animals were fasted for 2 h before IP injection with insulin (Humulin-R, Eli Lilly) at 0.75 U/kg in in 0.9% saline. The insulin was prepared from a 100 U/mL stock vial and was administered using a 27-gauge needle attached to 1-mL syringe. The animals are weighed before testing to determine the correct dose, and the blood glucose was measured as described above at times 0 (before insulin injection) and then after 15, 30, 45 and 60 min. For each mouse, the blood glucose levels at each time point were calculated as percentage of the blood glucose levels at time 0. Mean glucose values (± SEM) for each strain were plotted against time. Considering insulin losses in the tubes and the syringe, and the first passage through the liver with a 1% absorption and a total blood volume of 2 mL, the insulin concentration would theoretically be ∼125 µU/mL (or 5 ng/mL), which is in the supra-physiological range (3–5 fold after feeding).

### IHC staining method

Tissues for IHC were prepared by submerging dissected organs in cold 4% paraformaldehyde (PFA) in PBS and incubating at 4°C for 3–24 h. The tissues were then rinsed (2 × 15 min) in cold PBS at room temperature before transferring to 30% sucrose in PBS and incubating overnight at 4°C. The sucrose was then replaced with a 1:1 mixture of 30% sucrose in PBS and OCT, and incubated for 15 min rocking at room temperature. The mixture was replaced with 100% OCT and incubated as above. The organs were then transferred to label molds, covered with OCT and placed on dry ice until opaque and hard. A Cryostat CM1950 was used to prepare 8-μm sections on permafrost Fisher slides (AML Laboratories), which were stored at –80°C in a covered box. Multiple slides (n=20) were obtained for pancreas, spleen, liver, white adipose, liver, brown adipose, brain and quad muscle samples for each mouse.

The sections were stained by circling the tissue with a hydrophobic pen and adding 200 μL 4% PFA in PBS for 15 min at room temperature. The slides were then rinsed in PBS, incubated in 0.3% H_2_O_2_ in PBS for 15 min at room temperature, and rinsed again in PBS, before permeabilization in 0.1% Triton-X 100 in PBS for 15 min at room temperature. The slides were then rinsed in PBS, air dried, and blocked with 200 μL 0.5% BSA and 1% Tween-20 in PBS for 30 min at room temperature. After air drying, the sections were incubated with a primary mouse mAb specific for AdipoR1-CTF (clone 444) conjugated to FITC (0.5 μg/mL in PBS) along with a guinea pig mAb specific for insulin (Pierce, PA26938) diluted 1:100 or a rat mAb specific for CD45 (Thermo Fisher Scientific, MA1-70096) diluted 1:200. The slides were incubated overnight at 4°C before washing with 0.1% Tween-20 in PBS for 5 min followed by PBS for 2 min. We then added the secondary antibodies (anti-guinea pig conjugated to Alexa 647, donkey pAb Jackson ImmunoResearch, 706-605-148; or anti-rat conjugated to Alexa 647, donkey pAb Jackson ImmunoResearch, 712-606-153), each diluted to 2.5 μL/mL in PBS, and incubated the slides at room temperature for 1 h before washing as above. The slides were counterstained with 50 μL 1 μg/mL 4,6 diamidino-2-phenylindole dihydrochloride (DAPI) followed by rinsing in PBS. After mounting a cover slip with 5 μL DAPCO, phase contrast and fluorescence microscopy was conducted with a Zeiss Axioscan using Zen Blue and appropriate filter sets for DAPI, FITC and Alexa 647. The slides were scanned at 40× using 50–300 ms exposures. A grading system was established for consistency across all slides in an experiment.

### Measurement of β cell mass and insulitis scoring

Pancreata were harvested, embedded in paraffin, and sectioned, before staining for insulin using an insulin-specific antibody. The β cell mass and insulitis scores were determined on three slides collected throughout the pancreas.^10,13,15^ For insulitis scoring, all islets in a given pancreas section were scored blindly by two people using the following scale: Grade 1 = no islet-associated mononuclear cell infiltrates; Grade 2 = peri-insulitis affecting <50% of the circumference of the islets without evidence of islet invasion; Grade 3 = peri-insulitis affecting >50% of the circumference of the islets without evidence of islet invasion; Grade 4 = islet invasion.

### Islet protocol

Islets prepared by the IUMS islet core were picked after resting overnight at 37°C in a 5% CO_2_ atmosphere. Picked islets were centrifuged at 1000 × *g* for 5 min and the supernatant was discarded. The remaining islet pellet was resuspended in 60 µL lysis buffer (50 mM Tris pH 8.0, 150 mM NaCl, 0.05% deoxycholate, 0.1% IGEPAL (NP40), 0.1% SDS, 0.2% sarcosyl, 10% glycerol, 1 mM DTT, 1 mM EDTA, 10 mM NaF, protease inhibitors (complete Mini, EDTA-free, Roche), phosphatase inhibitors (PhosStop, Roche), 2 mM MgCl_2_, 0.05% benzonase) via trituration and stored at –80°C. At the end of the study, samples were thawed and 5 µL was used for a Pierce BCA assay (Thermo Fisher Scientific). Samples were diluted to 0.5 mg/mL in 6x Laemmli sample buffer and 5 µg of total islet protein was used for SDS-PAGE and western blotting.

### Mouse tissue protocol

Tissue samples were stored at –80°C until the end of the study, then spleen and pancreas samples were suspended in 1 mL lysis buffer (see above) and homogenized in 5-mL Eppendorf microfuge tubes. After centrifugation at 1000 × *g* for 10 min at 4°C, 800 µL of the supernatant was transferred to a fresh microfuge tube and centrifuged at 5000 × *g* for 5 min at 4°C. Then, 750 µL of the remaining supernatant was centrifuged at 6000 × *g* for 5 min at 4°C. Finally, 500 µL of the remaining supernatant was transferred to a fresh microfuge tube and 10 µL was used for a BCA assay, while the remaining sample was stored at –80°C. Based on the BCA assay results, the samples were thawed on ice and diluted to 4 µg/mL in 6x Laemmli sample buffer. We then used 40 µg of mouse tissue lysate for western blotting.

### Western blotting protocol

We loaded 4–20% TGX gels (Bio-Rad) with tissue lysate and 1–2 µL of the Chameleon 800 ladder (LI-COR). The samples were fractionated in 1× Tris/glycine/SDS buffer (Bio-Rad) for 90 minutes at 100 V. The samples were transferred to LF-PVDF membranes (Bio-Rad) in 1× Tris/glycine buffer with 20% methanol at 100 V for 60 min. Membranes were blocked with Odyssey blocking buffer (LI-COR), and probed with the appropriate antibodies overnight at 4°C. After rinsing, the membranes were incubated for 1 h at room temperature on an orbital shaker with IR800 and IR680RD secondary antibody conjugates (LI-COR) diluted 1:20,000 in Odyssey blocking buffer containing 0.01% SDS. The membranes were washed 3 × 5 min in 10 mL Tris-buffered saline (TBS) containing 0.05% Tween-20 and then 1 × 5 min in TBS before image acquisition on an Odyssey Clx scanner.

### Splenocyte isolation method

Euthanized mice were placed on a clean dissection board and rinsed with 70% alcohol. An incision was made in the abdominal cavity, and the spleen was dissected and transferred to 2 mL cold RPMI medium. The cells were washed through a sterile 100-μm cell strainer into a 50-mL conical tube using 5 mL of RPMI medium. The cell suspension was centrifuged at 450 × *g* for 6 min at 4°C, the supernatant was removed, and the cell pellet was resuspended in 2 mL RBC lysis buffer (Thermo Fisher Scientific). The cells were incubated for 3 min at room temperature and mixed with 10 mL PBS before centrifuging as above and removing the supernatant. The final pellet was resuspended in 5 mL RPMI medium supplemented with 10% FBS, 100 Units/ml of penicillin/streptomycin and 10 mM HEPES. The cells were counted using a hemocytometer and viability was determined by trypan blue staining.

### FACS analysis

Pancreatic lymph nodes (pLNs) and spleens were processed for T-cell counting (Th1, Th17 and T_reg_ populations) as previously described.^13,14^ Briefly, pLN cells and splenocytes freshly isolated from C57 and NOD mice were stained with fluorochrome-conjugated antibodies specific for CD4 (RM4-5, Biolegend), FoxP3 (MF23, BD Pharmigen), IFN-γ (XMG1.2, BD Pharmigen) or IL-17 (TC11-18H10). For intracellular staining, single cells were stimulated for 4 h with 100 ng/mL PMA (Sigma-Aldrich), 500 ng/mL ionomycin (Sigma-Aldrich), and GolgiPlug (BD), at 37°C in a 5% CO_2_ atmosphere. Cells were stained with anti-CD4, then fixed and permeabilized in a combination buffer (Themo Fisher Scientific). Intracellular antibodies were added and cells were incubated for 30 min at 4°C in the dark. After staining, the cells were washed and analyzed in a FACSCanto II cytometer (BD Pharmigen). Data were analyzed using FlowJo software (Tree Star). Initially, the cells were gated on FSC-A and FSC-H for doublet exclusion, followed by gating on FSC (size) and SSC (granularity) to select a population compatible with lymphocytes, within which the CD4^+^ CD25^+^ Foxp3^+^ CD4^+^ IFN-γ^+^ or CD4^+^ IL-17 ^+^ cells were evaluated.

### Blood analysis

Serum samples were tested for the presence of CTF and CTF-specific autoantibodies (IgG-CTF) using sandwich ELISAs as previously described,^9^ although the ALP-conjugated anti-human IgG antibody was replaced with an anti-mouse equivalent (Sigma A3562). Blood samples were collected from all animals 14 days prior to treatment and also prior to the GTT/IPITT procedures (104 days) and were centrifuged to collect 30-μL aliquots of serum. Samples for insulin ELISA (2 × 5 μL duplicates) were stored in 96-well break-a-way plate strips with individual flat caps (Phenix #MPC-4896 and #MPC-0100). Insulin was measured using the Alpco Stellux ELISA kit for rodent samples. For IgG-CTF testing, 10-μL samples were diluted to 480 μL with a stabilization buffer comprising 1% BSA (1 pkg/L, Sigma-Aldrich), 0.1 M citrate (ACS grade monohydrate, Sigma-Aldrich) and 0.027 g/L EDTA sodium salt (Sigma-Aldrich) in PBS (pH 6.4). For free CTF testing, 10-μL samples were diluted to 120 μL with stabilization buffer comprising 25 mM Tris and 0.15 M NaCl with Pierce Protease Inhibitor Mini Tablets (two tablets in 20 mL buffer). Terminal blood samples were collected from all mice, and serum samples were prepared for the analysis of free CTF, IgG-CTF and insulin, with an additional 40 μL of serum mixed with 480 μL Ig-CTF buffer for western blot analysis. Samples were prepared in duplicate wells in a polypropylene sample plate. The sample plate was sealed tightly with foil and stored at 4°C for testing within 24 h or stored at –70°C for up to 1 year for repeat testing. Mouse blood was tested for adiponectin by ELISA (Thermo Fisher Scientific KRP0041).

## Results and discussion

### Complexes containing mouse IgG and CTF accumulate in diabetes

In our first study, we found that the abundance of IgG-CTF complexes in the serum increased with age in male C57 and female NOD mice (Figure 1A; & **Supplemental Report 1**). The abundance of free CTF was significantly higher in female NOD mice than in male C57 mice prior to the onset of an autoimmune response (Figure 1B).

**Figure 1:**
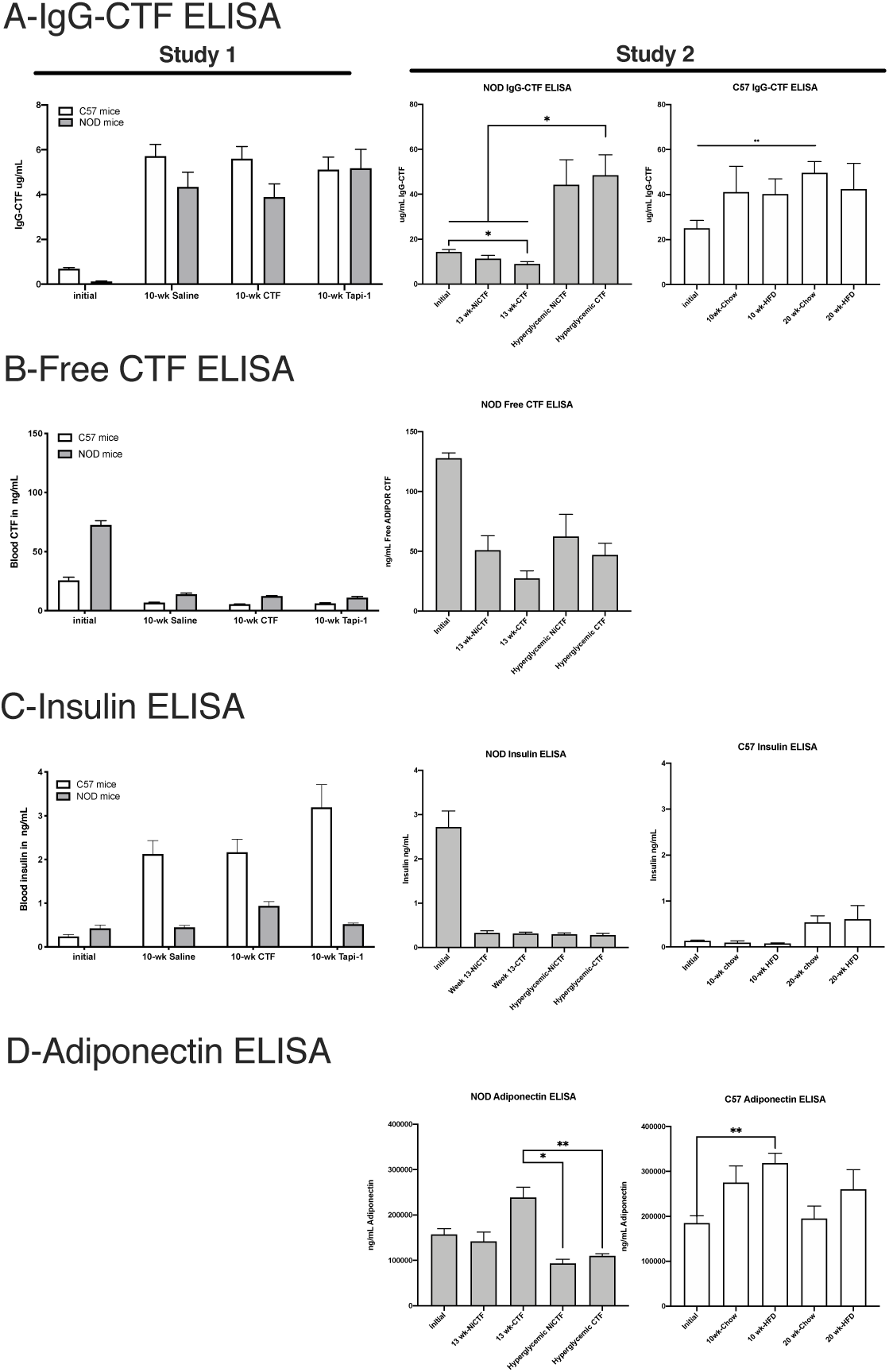
Plasma ELISA measurements for IgG-CTF, CTF, insulin, and adiponectin. Analysis of mouse sera by ELISA to detect the IgG-CTF complex (A), free AdipoR1-CTF (B), insulin (C), and adiponectin (D) (n = 5–70 per group). Initial measurements were taken at 0 weeks in the study, and final measurements were taken following two fed blood glucose measurements of >250mg/dL (hyperglycemic group) or 13 weeks after peptide dosing. Data shown means ± SEM (*values are significantly different from other columns by one-way ANOVA, 0.05>P).

In our second study, we found that the abundance of IgG-CTF complexes also increased significantly after 20 weeks on a normal diet in female C57 mice and with age in female NOD mice (Figure 1A). The abundance of free CTF was again found to rise prior to the onset of an autoimmune response and to diminish as autoantibodies formed, but the amount of unbound endogenous CTF did not correlate with the onset of diabetes nor was it significantly increased by exogenous CTF treatment (Figure 1B). We anticipated that exogenous CTF would not be exchanged *in vivo* because autoantibody complexes do not release endogenous CTF other than under strongly basic conditions, and did not exchange with exogenous synthetic CTF *in vitro*.

The second study also showed that free CTF in the C57 mice was below the limit of detection. Interestingly, insulin levels were higher in NOD mice before the onset of an autoimmune response, decreasing with onset, and were lower in DIO mice prior to the start of the high-fat diet (Figure 1C). This was consistent with the data from the NOD mice in the first study. Insulin levels increased with obesity in C57 mice, but not as rapidly as in the study demonstrating hyperinsulinemia after 10 weeks on the high-fat diet (Figure 1C). Notably hyperglycemia correlated with a decrease in serum adiponectin compared to non-hyperglycemic NOD mice (Figure 1D). The high-fat diet caused a significant increase in serum adiponectin levels in C57 mice (Figure 1D).

### CTF peptide treatment increases insulin sensitivity in NOD mice

Weekly dosing of NOD mice with 1 mg/kg exogenous CTF peptide containing the IDE-binding domain caused a slight but significant increase in insulin sensitivity (IPITT) after 8 weeks of treatment compared to the NiCTF peptide lacking the IDE-binding domain Figure 2B). This is consistent with our proposal that the inhibition of circulating IDE could reduce insulin degradation in the plasma, and suggests that previously observed increases in insulin levels 120 min after exogenous CTF injection probably reflect the short-term inhibition of circulating IDE in the peripheral blood. There was no significant difference between the peptide treatments in terms of glucose tolerance, weekly blood glucose, weekly body weight, or conversion to diabetes (Figure 2A, 2C, 2D). Furthermore, neither of the exogenous CTF peptides affected the IgG-CTF levels in plasma, body weight or conversion of NOD mice to diabetes. A supplemental experiment showed that the AdipoR1-CTF peptide did not enter cells (macrophages) under viable growth conditions and did not affect intracellular insulin levels (**Supplemental Report 3**).

**Figure 2:**
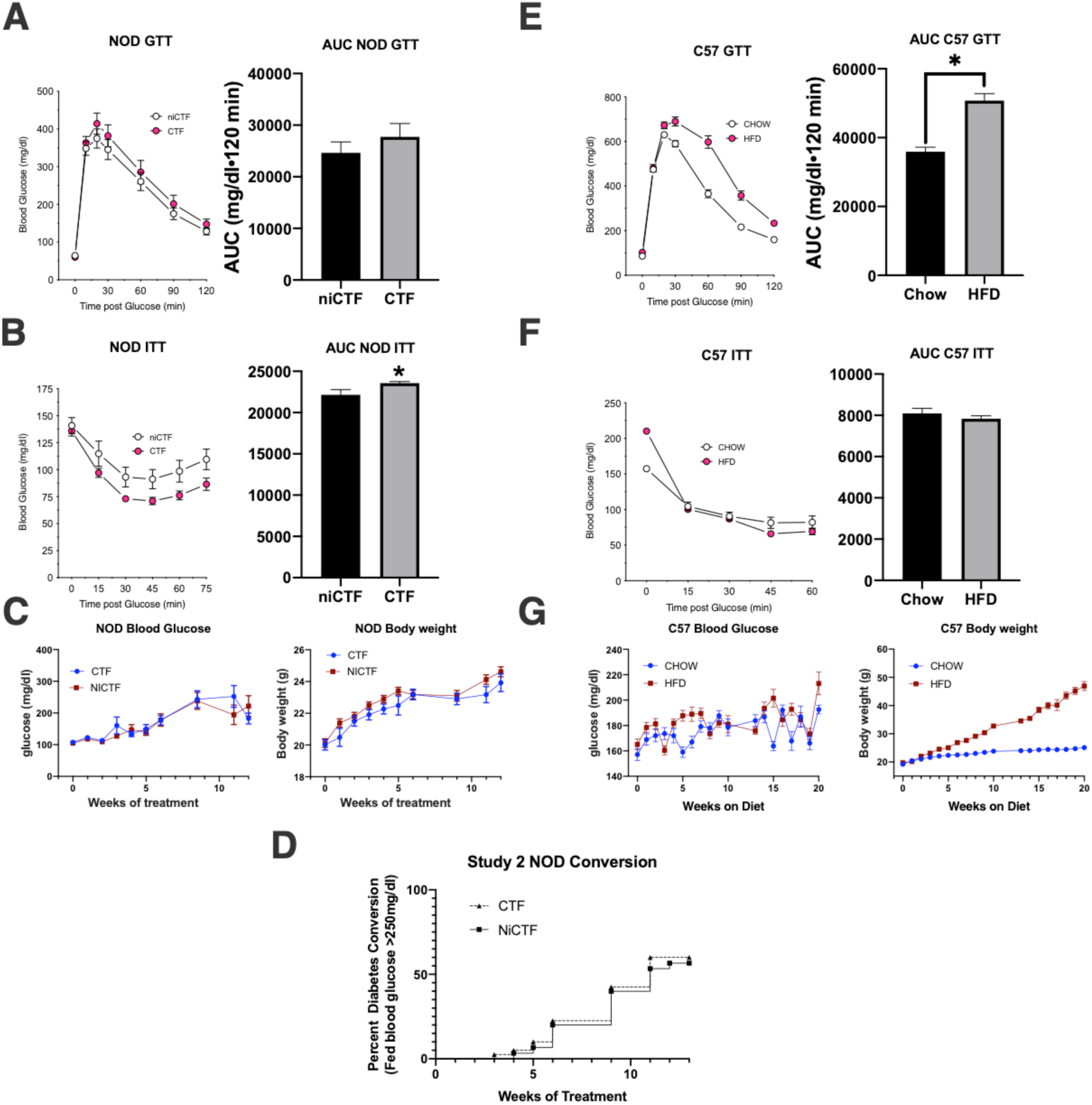
NOD and C57 intraperitoneal insulin and glucose tolerance tests (IPITT, IPGTT) showing mouse body weight, glucose, and NOD incidence of hyperglycemia. NOD mice dosed weekly with CTF or NiCTF (A-C) and C57 mice on normal or high-fat diets (E-G) were tested by IPGTT after 7 weeks of weekly peptide dosing (A) and 11 weeks of diet (E), and were tested by IPITT after 8 weeks of peptide dosing (B) and 12 weeks of diet (F). IPITT and IPGTT curves are accompanied by measurements of the area under the curve to determine statistical significance. Data show means ± SEM (*values are significantly different from other columns in an unpaired t-test, 0.05>P). Blood glucose and body weight were measured weekly except for the weeks that the IPITT and IPGTT tests were performed (C and G). The percent of the NOD cohort that were classified as diabetic was included on a weekly basis (D).

### Intact adiponectin receptor levels decrease in diabetic NOD mice but not in C57 mice

Western blots of pancreas lysate were prepared from mice euthanized at the start of the study and those euthanized due to hyperglycemia, revealing that the abundance of the adiponectin receptor decreased significantly in hyperglycemic mice independent of the peptide treatment (Figure 3). The levels of intact receptor in the pancreas were consistent with preliminary comparisons by ICC using anti-CTF antibodies (**Supplemental Report 1**). Differences in the level of intact receptor between the mouse models were observed in the pancreas but not in the liver, brown adipose, muscle or spleen tissues. Additional western blot analysis of adiponectin receptor levels in the pancreas, islet and spleen tissue from a 4-week-old C57 mouse confirmed that the intact receptor is expressed at much higher levels in the exocrine pancreas than the islets (**Supplemental Report 1**). Pancreatic lysate was prepared from 6-week-old C57 mice and those fed for 10 weeks on a normal or high-fat diet. Depending on the antibody probe, western blots showed that adiponectin receptor expression increased in mice fed on a normal diet (mouse mAb 444), or those fed on the high-fat diet (rabbit mAb against the receptor C-terminus) or that there was no significant difference (rabbit mAb against the receptor N-terminus). These data show that receptor expression is either unchanged or slightly higher after 10 weeks on either diet (Figure 4). We also found that adiponectin receptor expression was unchanged in the spleen lysate independent of age, peptide treatment or diet in both mouse models (Figure 5).

**Figure 3:**
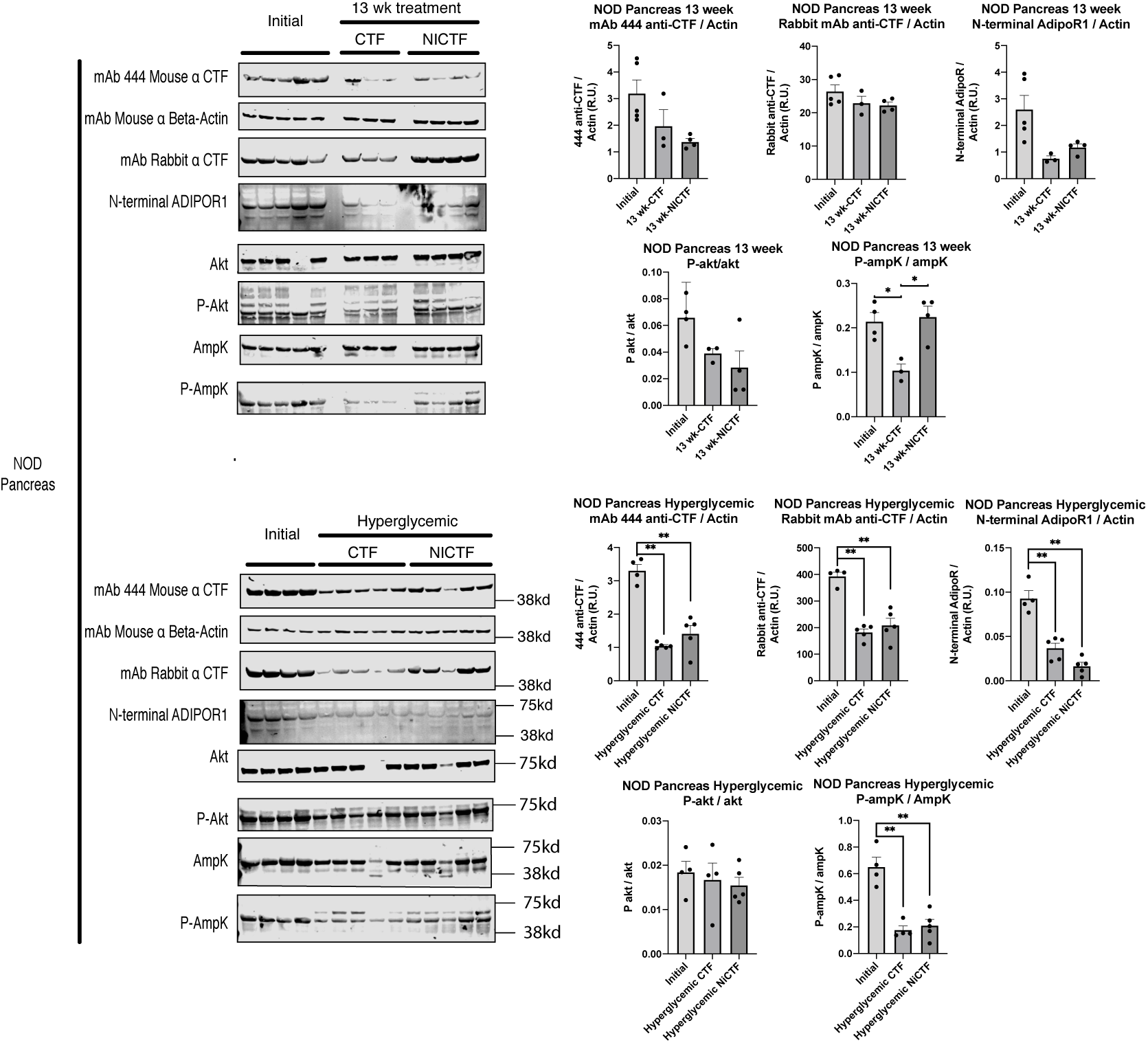
Adiponectin receptor expression with Akt and AmpK activation in NOD pancreata. NOD mice were euthanized according to study protocol, and pancreas tissue lysates were tested by western blot (40 µg of protein per lane) using the indicated antibodies. Quantitation was performed using Image Studio Lite. Adiponectin receptor antibodies were normalized against β-actin. P-Akt and P-AmpK antibodies were normalized using Akt and AmpK expression, respectively. Data show means ± SEM (*values are significantly different from other columns by one-way ANOVA, 0.05>P).

**Figure 4:**
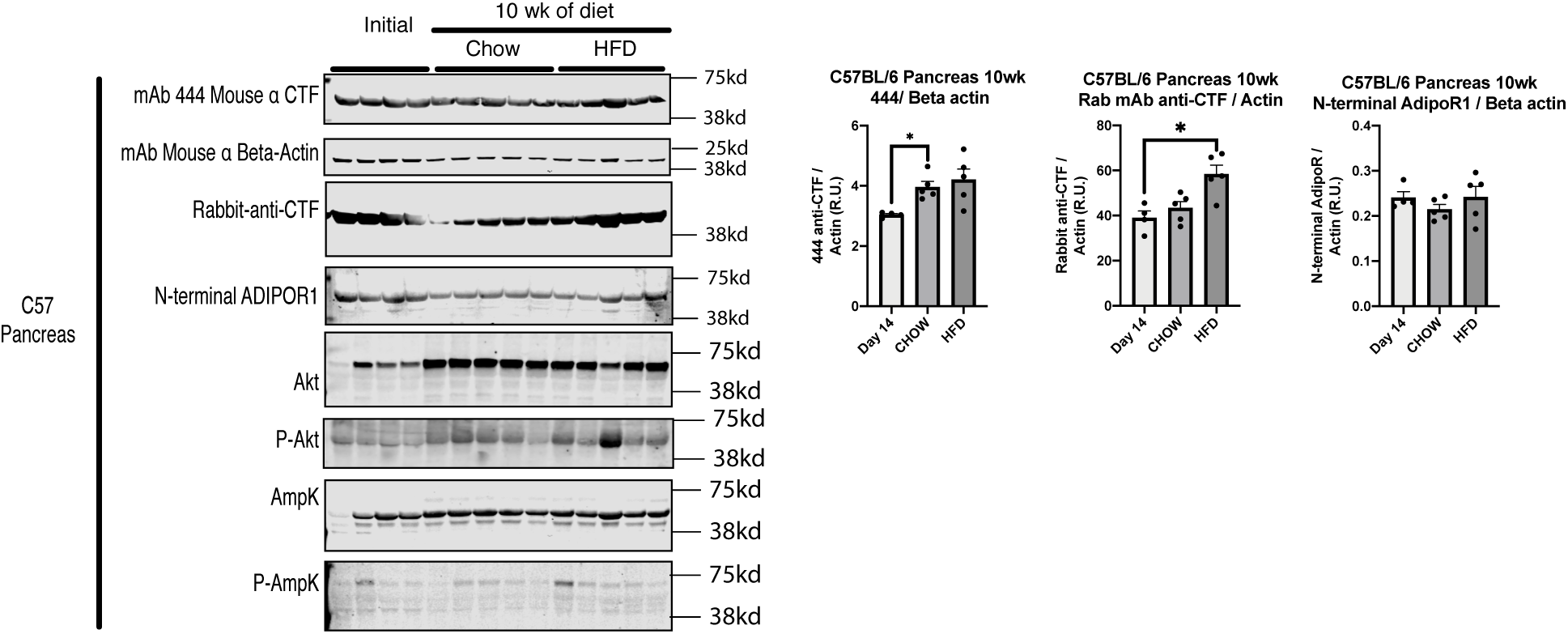
Adiponectin receptor expression with Akt and AmpK activation in C57 pancreata. C57 mice were euthanized according to study protocol, and pancreas tissue lysates were tested by western blot (40 µg of protein per lane) using the indicated antibodies. Quantitation was performed using Image Studio Lite. Adiponectin receptor antibodies were normalized against β-actin. Data show means ± SEM (*values are significantly different from other columns by one-way ANOVA, 0.05>P).

**Figure 5:**
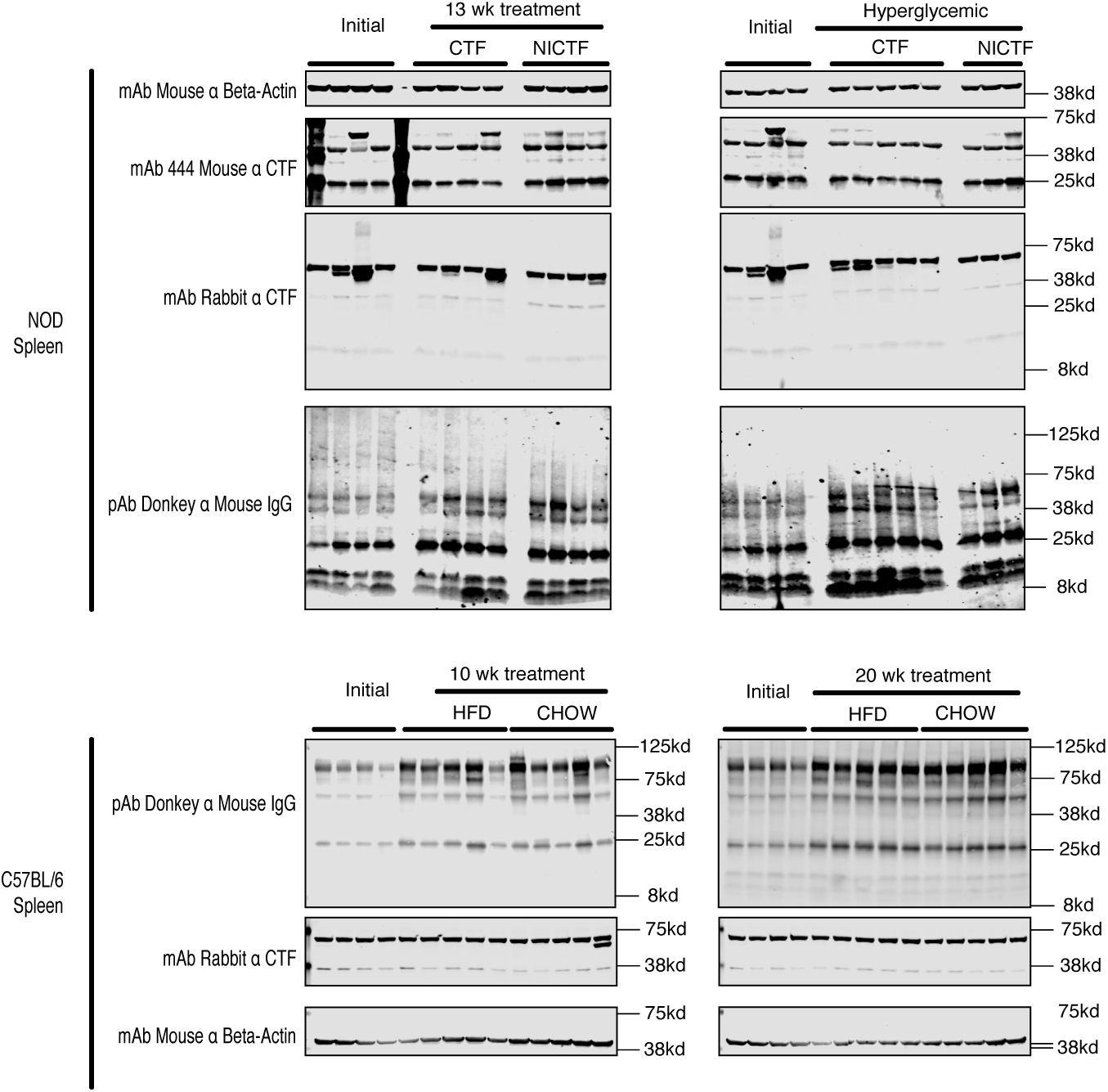
Adiponectin receptor expression in spleen lysates from NOD and C57 mice. The mice were euthanized according to scheme 1 and spleen tissue lysates (40 µg per lane) were tested by western blot (40 µg of protein per lane) using the indicated antibodies.

This loss of intact receptor in the NOD model was matched by a significant decrease in AMPK activation in hyperglycemic NOD mice without a significant change in AKT activation, in both cases independent of peptide treatment (Figure 3). There was no significant difference in receptor expression in non-hyperglycemic NOD mice (Figure 3). Treatment with the exogenous CTF lacking the IDE-binding domain (NiCTF) correlated with significantly higher AMPK activation compared to mice euthanized at the start of the study and age-matched CTF-treated mice, whereas AKT activation was not significantly dependent on age or peptide treatment (Figure 3). The effect of the IDE-binding domain present in the exogenous CTF was only pronounced in the non-hyperglycemic NOD mice, whereas no effect was observed in hyperglycemic NOD mice.

### CTF peptide treatment does not prevent insulitis

NOD mice were euthanized on day 105 of treatment or following the onset of hyperglycemia, and pancreata were harvested for insulitis scoring. Three slides were analyzed per pancreas and the total number of islets was counted and scored for insulitis (Figure 6). The total number of islets decreased with age, and decreased significantly with hyperglycemia independent of peptide treatment. Average insulitis scores significantly increased with age and hyperglycemia, independent of peptide treatment (Figure 6).

**Figure 6:**
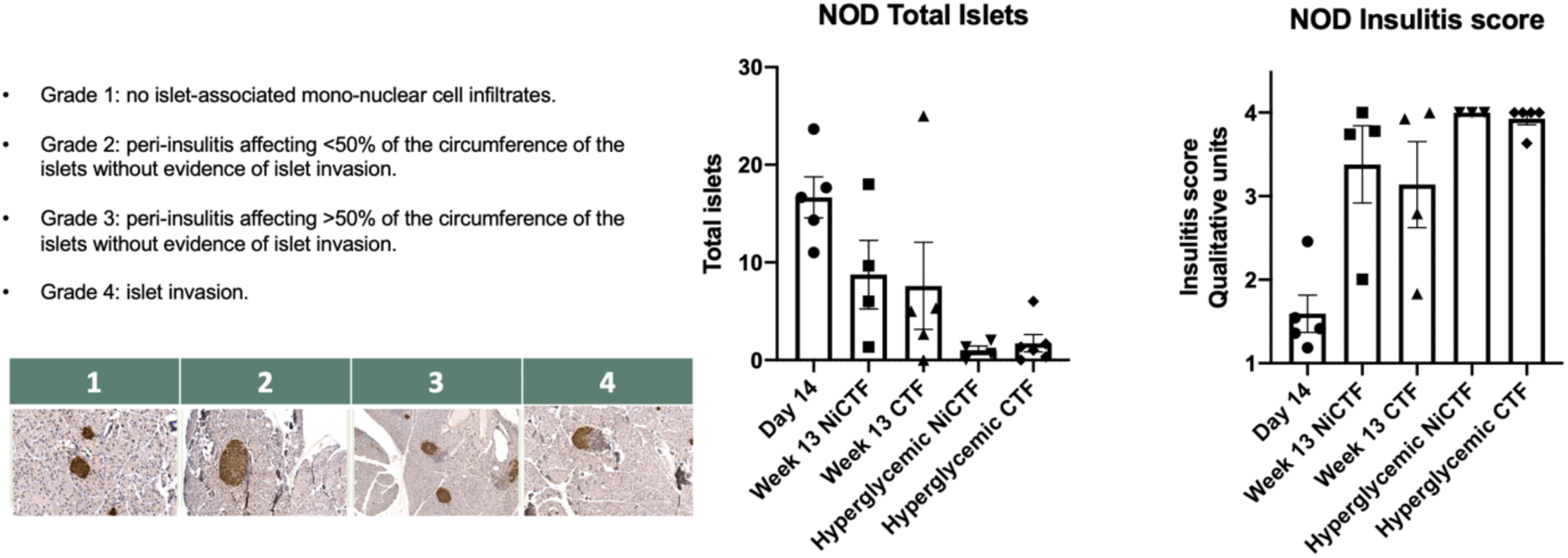
NOD insulitis scoring. NOD pancreata were collected as noted in scheme 1 and embedded in paraffin. Sections were stained for insulin, and islets were enumerated and scored for insulitis according to the grades shown.

### Changes in T_reg_, Th17 and Th1 cell populations in NOD and DIO mice

We compared the populations of T_reg_ (CD4^+^CD25^+^Foxp3^+^), Th1 (CD4^+^IFNγ^+^) and Th17 (CD4^+^ IL17a^+^) cells in the pancreatic lymph nodes (pLNs) and spleens of NOD and DIO mice by flow cytometry. In NOD mice, the number of T_reg_ cells in the spleen increased with age independent of hyperglycemia and peptide treatment, but in the pLNs the T_reg_ population increased with age, but not hyperglycemia (Figure 7). In DIO mice, the number of T_reg_ cells in the spleen increased with age but not in pLNs. The Th1 population in the spleen and pLNs of NOD mice increased significantly with age, but not hyperglycemia (Figure 7). The number of Th17 cells in the spleen and pLNs of NOD mice increased significantly with hyperglycemia compared to initial values and to mice treated for 13 weeks with NiCTF (Figure 7).

**Figure 7.**
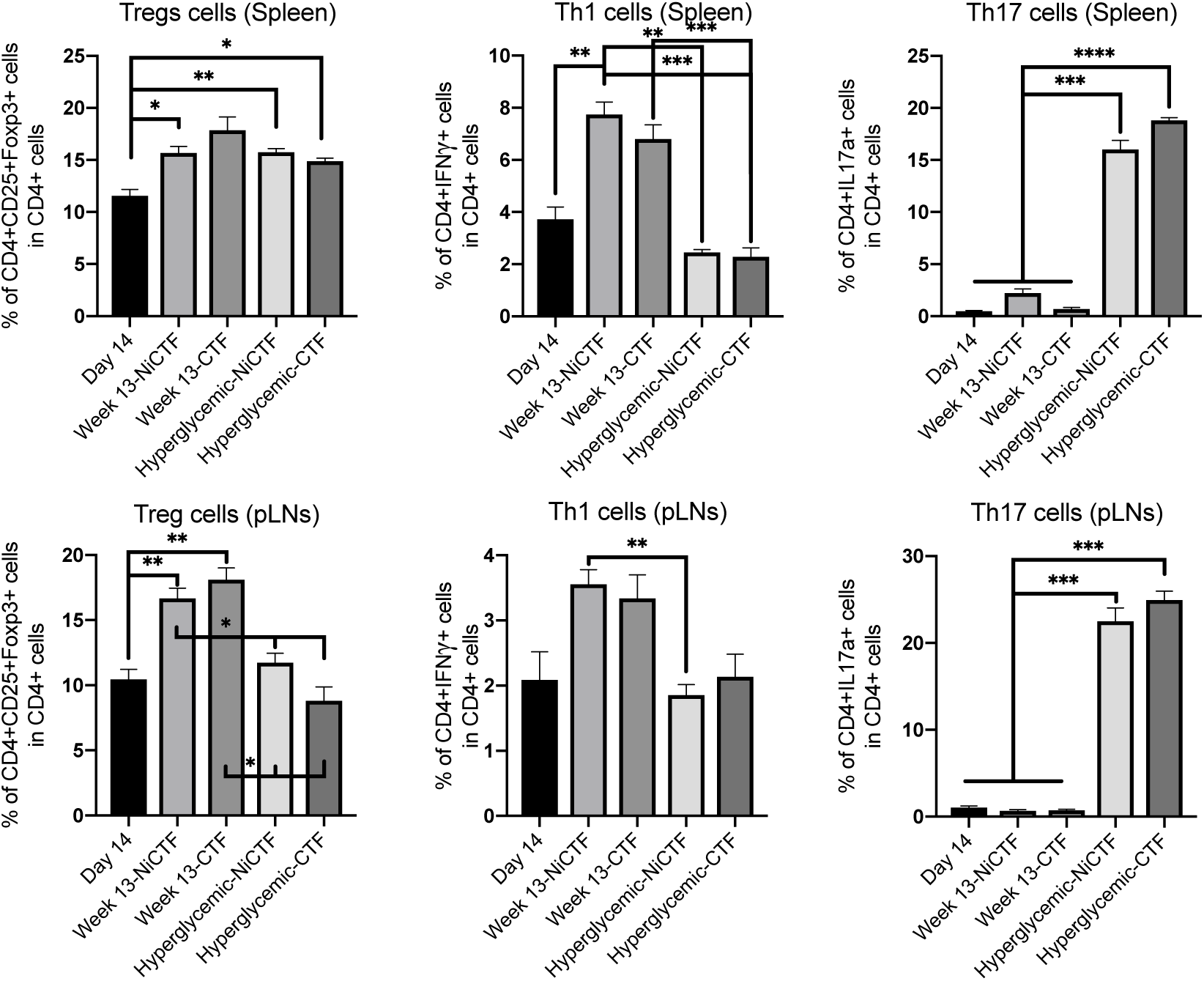
CD4^+^ T-cell populations the spleen and pLNs of NOD mice dosed with the NiCTF or CTF peptides according to scheme 1. Following the onset of diabetes or upon reaching 13 weeks of age, the Th1, Th17 and T_reg_ cell populations were characterized by flow cytometry (n = 3-5 per group). Data show means ± SEM (*values are significantly different from other columns by one-way ANOVA).

In C57 mice, the T_reg_ cell population significantly increased with age in the spleen but not in the pLNs (Figure 8). The number of Th1 cells in the spleen and pLNs significantly increased after 10 weeks on the high-fat diet, but between 10 and 20 weeks on either diet there was a decline in the number of Th1 cells in the spleen and no change in the pLNs. The number of Th17 cells in the spleen significantly increased after 20 weeks on the normal or high-fat diet, and in the pLNs the number of Th17 cells increased after 20 weeks on the normal diet (Figure 8).

**Figure 8:**
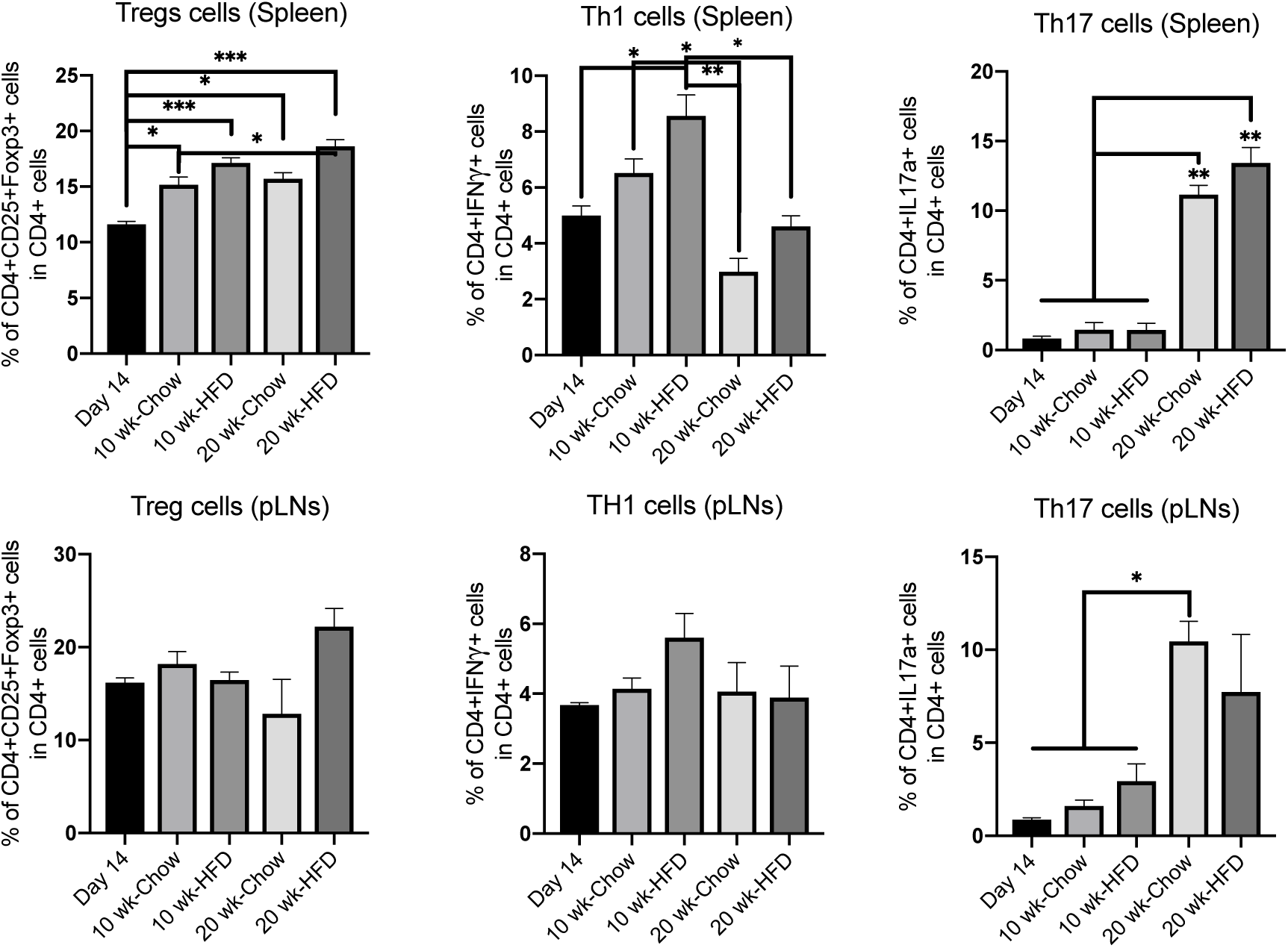
CD4^+^ T-cell populations the spleen and pLNs of C57 mice on a high-fat diet compared to controls on a normal diet. At 16 and 26 weeks of age (10 and 20 weeks on diet), the Th1, Th17 and T_reg_ cell populations were characterized by flow cytometry (n = 3-5 per group). Data show means ± SEM (*values are significantly different from other columns by one-way ANOVA).

Although an inflammatory response was observed in both the obese and non-obese models, it was more pronounced in the non-obese model, which progresses to islet cell death and autoimmune disease. There was an increase in the number of Th17 cells in both models in the absence of progression, and an increase in the number of Th1 cells in both models during progression. The changes were more pronounced in the non-obese model, which also showed a more significant increase in the T_reg_ cell population in the absence of progression.

## Discussion

Previous reports have shown human islets and rodent β cells express *AdipoR1* and *AdipoR2* mRNA at levels greater than in muscle and brain, and comparable to the levels in liver.^16^ Furthermore, *AdipoR1* and *AdipoR2* gene expression decreases by ∼40% in the pancreas of obese mice relative to lean littermates, and AdipoR1 and AdipoR2 predominantly accumulate in the acinar cells of the exocrine pancreas.^17^

We found that adiponectin receptor expression decreases significantly in hyperglycemic NOD mice, with a concurrent loss of AMBK signaling. In the C57 model of DIO, the abundance of the receptor increased with age and in mice fed on a high-fat diet, and AMPK signaling was preserved. We observed increases in the numbers of inflammatory T_reg_, Th1 and Th17 cells in the spleen and pLNs that correlated with age, hyperglycemia, and diet in our studies of NOD mice (T1D model) and C57 mice on the high-fat diet (T2D model). The number of Th17 cells increased most significantly in NOD mice progressing to hyperglycemia.

Many inflammatory genes and pathways are upregulated in T1D, including the expression of proteases, so it is difficult to pinpoint the agent responsible for the proteolytic degradation of the adiponectin receptor.^18^ We found that the degradation of the receptor was not directly related to its expression levels. Although TACE is expressed at significantly higher levels in the NOD model, the administration of a TACE inhibitor did not prevent CTF formation, suggesting additional factors are involved in this process.

## Conclusion

We observed the more pronounced shedding of CTF from the adiponectin receptor into the peripheral blood in the TACE-active NOD strain (T1D model) which represents an inflammatory autoimmune phenotype, but fragmentation was also observed to a lesser extent in the C57 strain with DIO (T2D model). Autoantibodies to CTF were observed in both models. Neither of the exogenous CTF peptides had impact on the IgG-CTF complex levels in plasma, body weight or the conversion of NOD mice to diabetes. In summary, we describe a pattern of adiponectin receptor fragmentation and autoantibody production under physiological conditions of aging, diet-inducted obesity, and autoimmune diabetes.

## Supporting information

Supplement Report 1. Murine study of AdipoR CTF formation

Supplement Report 2. Antibody blocking of AdipoR CTF inhibition of IDE

Supplement Report 3. Macrophage study of AdipoR CTF Formation

## Acknowledgements

The Indiana Bioscience Research Institute (an independent non-profit research organization) sponsored the laboratory work behind this study.

## ABBREVIATIONS

ACC: Acetyl coenzyme A carboxylase
AKT: Protein kinase B
AMPK: 5’-AMP-activated protein kinase
CTF: C-terminal fragment of the adiponectin receptor GLUT Glucose transporter
GTT: Glucose tolerance test
IDE: Insulin degradation enzyme
ITT: Insulin tolerance test
IP: Intraperitoneal
MAPK: Mitogen-activated protein kinase
PPAR: Peroxisome proliferator-activated receptor T1D Type 1 diabetes
T2D: Type 2 diabetes
TACE: Tumor necrosis factor-α cleavage enzyme (aka ADAM-17, MMP-17)
TNFα: Tumor necrosis factor-α

**Supplement Report 1**. Formation of the adiponectin receptor C-terminal fragment by TACE in mice

**Supplement Report 2.** Blocking the inhibition of IDE by the C-terminal fragment of the adiponectin receptor using a specific autoantibody

**Supplement Report 3.** The influence of macrophage differentiation and polarization on the formation of the adiponectin receptor C-terminal fragment

**Conclusion of study 1:** The cause of adiponectin receptor fragmentation was not identified in this study, but pancreatic tissue in mice know to have higher TACE activity, also have greater AdipoR1 fragmentation.

**Conclusion of study 2:** This study supports the neutralization of plasma IDE inhibition by CTF autoantibodies. We found that antibodies specific for the AdipoR1-CTF_351-362_ domain were able to block the non-competitive inhibition of IDE by CTF.

**Conclusion of study 3:** Viable human blood cells in culture did not take up exogenous CTF, and no direct impact on intracellular insulin was observed. TACE activity did not explain the endogenous fragmentation of the receptor, but we observed a correlation with TNFα release and endogenous fragmentation occurred following the polarization of M0 to M1 macrophages.

