## Supplement Report 1. Murine study of AdipoR CTF formation for "Adiponectin Receptor Fragmentation in Mouse Models of Type 1 and Type 2 Diabetes"

### Formation of the adiponectin receptor C-terminal fragment by TACE in mice

##### Introduction

The C-terminal fragment of the adiponectin receptor (AdipoR-CTF) was previously observed to accumulate in the plasma during progression to diabetes in Zucker Diabetic Sprague Dawley (ZDSD) rats.[1] The quantity of endogenous AdipoR-$CTF_{344\text{-}375}$ in the plasma increased during the 120-min glucose tolerance challenge and was more pronounced in diabetic rats with worsening glucose tolerance. The levels of endogenous AdipoR1-CTF in the plasma were inversely correlated with the endogenous insulin plasma response. The administration of exogenous AdipoR1-$CTF_{351\text{-}370}$ to normal Sprague Dawley (SD) rats correlated with increased plasma insulin, but this was not the case in Zucker diabetic fatty (ZDF) rats with insulin insufficiency.[1] Autoantibodies against AdipoR1-CTF were detected in both diabetic ZDF and non-diabetic SD rats. Previous studies of enzyme inhibition revealed the competitive inhibition of tumor necrosis factor α cleavage enzyme (TACE) by AdipoR1-$CTF_{344\text{-}375}$, suggesting that TACE could be responsible for the production of endogenous CTF.[1] The endogenous and exogenous forms of AdipoR1-CTF are compared in Figure S1.

**Supplemental Figure 1 .** Endogenous AdipoR1 CTF 32-mer amino acid sequence highlighted and compared to synthetic exogenous AdipoR1 CTF 25-mer, and 18-mer peptides (displayed on right).

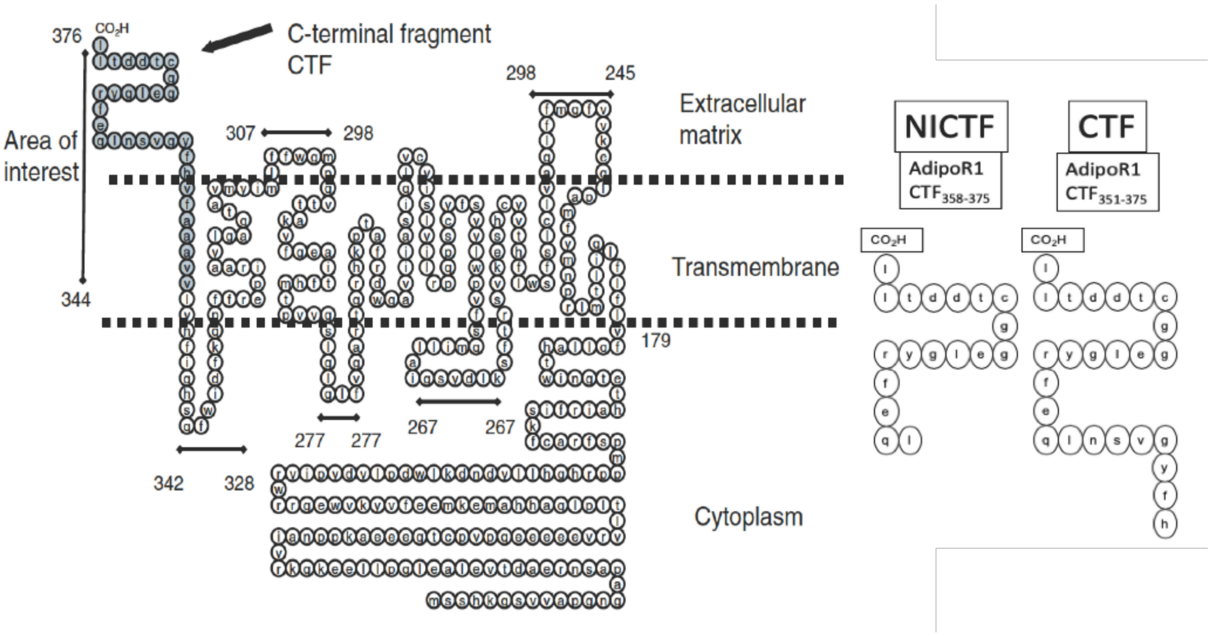

We were therefore motivated to explore the fragmentation of the adiponectin receptor and the resulting production of CTF in non-obese diabetic (NOD) mice, which provide an excellent model of insulin-dependent type 1 diabetes (T1D), and in C57 mice with diet-induced obesity (DIO), an excellent model of insulin-resistant type 2 diabetes (T2D). The NOD models show higher levels of TACE activity in the pancreas compared to DIO model.[2-4]

#### Experimental design and results

The NOD model was selected because 60–80% of females and 20–30% of males spontaneously develop diabetes at 16–30 weeks of age. The model is characterized by the autoimmune destruction of pancreatic islets, including elevated TACE expression, the migration of immune cells, and the onset of insulitis. These are advantageous characteristics to evaluate CTF release from the adiponectin receptor (Figure S2).

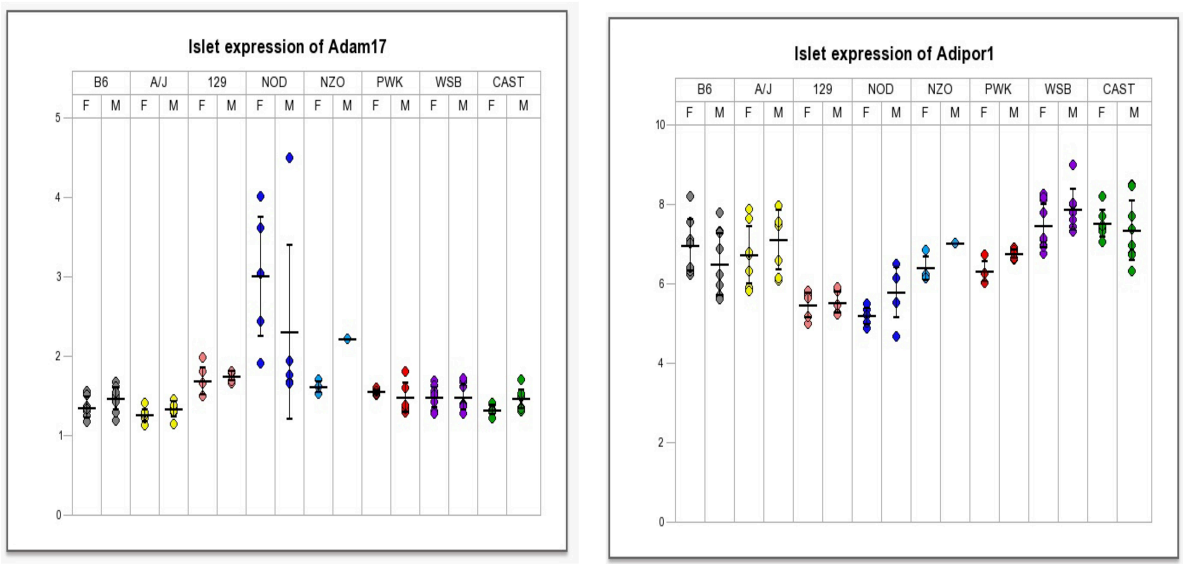


**Supplemental Figure 2. Expression of TACE (ADAM-17) and AdipoR1.** These panels summarize the islet RNA seq of mice models used for diabetes pathway research an http://diabetes.wisc.edu/cc_founder.php. **LEFT**: The TACE (ADAM-17) mRNA of islet of non-obese diabetic (NOD) are significantly elevated of Female over other models. **RIGHT**: ADIPOR1 mRNA is expressed in islet of non-obese diabetic (NOD) and all other models.

A strategy to monitor the formation of endogenous CTF and its autoantibodies during disease progression was developed, and was augmented by the administration of an exogenous AdipoR1-CTF$_{351-375}$ peptide and a TACE inhibitor (TAPI-1).[5] Animals were treated weekly with vehicle, AdipoR1-CTF$_{351-375}$ or TAPI-1 by intraperitoneal (IP) injections starting at 14 days (6 weeks of age) and continuing for up to 105 days (14 weeks of age). The design protocols of studies 1 and 2 are compared in Figure S3. The 16-week pilot study did not convert enough female NOD mice to diabetes to determine the statistical impact of CTF treatment on insulitis, and thus the study was expanded to 30 weeks (**Supplemental Report 2**).

**Supplemental Figure 3. Preliminary Study 1 protocol** for non-obese diabetic mice (NOD) and control mice (C57) on normal diet compare to Study 2 protocol non-obese diabetic mice (NOD) and control mice (C57) on high fat diet

In study 1, NOD and C57 mice were treated weekly either with vehicle, AdipoR1 $CTF_{351\text{-}375}$, or TACE inhibitor N-[(2R)-2-[2-(Hydroxyamino)-2-oxoethyl]-4-methyl-1-oxopentyl]-3-(2-naphthalenyl)-L-alanyl-N-(2-aminoethyl)-L-Alaninamide acetate salt (TAPI-1 acetate salt, Sigma-Aldrich, SML0739) and measured at A) 4-6 weeks of age prior to treatment, B) 16 weeks of age without treatment; C) 16 weeks of age with 10 week of synthetic TACE inhibitor treatment ; D) 16 weeks of age with 10 week of synthetic CTF treatment.

**Flow chart for final Study 2 protocol** is compared to the flow chart for preliminary study 1.

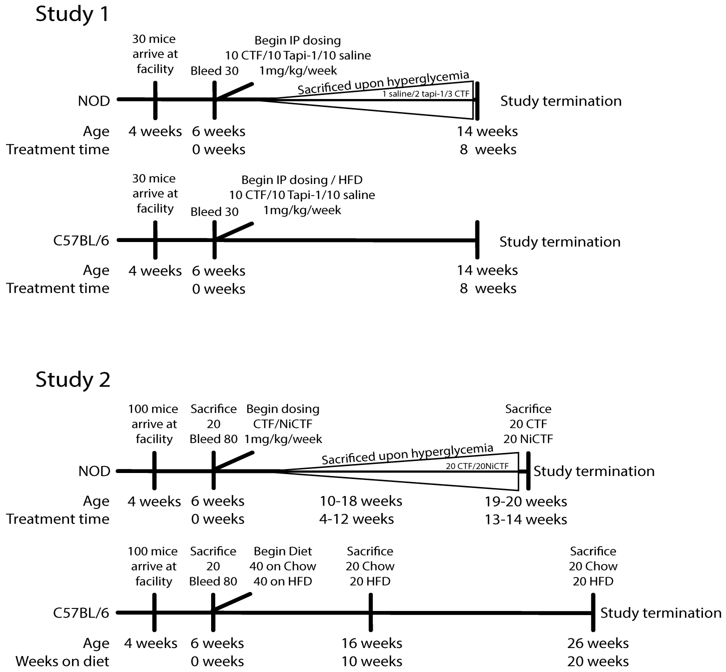


To determine whether the quantity of free CTF differs between the T1D and T2D models, we used a sandwich ELISA to compare blood samples from NOD and C57 DIO mice. We determined the quantity of free CTF and its autoantibodies (IgG-CTF) in blood samples taken at 4–6 weeks of age. In NOD mice, this is prior to a significant autoimmune response, and in C57 mice this is prior to the switch to a high-fat diet. We measured the levels again at 14 weeks of age, after 8 weeks of treatment with exogenous CTF or TAPI-1 (Figure S4).

The levels of free CTF were significantly higher in NOD mice than C57 mice at 4–6 weeks of age (Figure S4a). Both strains showed a significant reduction in the levels of exogenous free CTF as they aged to 16 weeks and progressed toward diabetes (Figure S4a). In contrast, the levels of IgG-CTF increased as the mice aged to 16 weeks (Figure S4b). The higher concentrations of free CTF did not correlate with lower blood glucose levels in samples collected from unfasted mice at 16 weeks of age (Figure S4c). The plasma insulin levels were higher in C57 mice after they were switched to the high-fat diet and were higher than the levels in NOD mice, as expected (Figure S4d).

Subsets of mice were then treated with TAPI-1 or synthetic CTF for 10 weeks to determine the effect on CTF and IgG-CTF levels, and blood samples were taken at 16 weeks (Figure S4a,b,d). CTF treatment had no effect on the level of plasma insulin. Furthermore, the TACE inhibitor did not influence the level of CTF in the plasma, suggesting that additional factors are involved in its formation. There was no significant difference in IgG-CTF levels between NOD and C57 mice, with both strains showing a rise in IgG-CTF levels between 6 and 16 weeks of age.

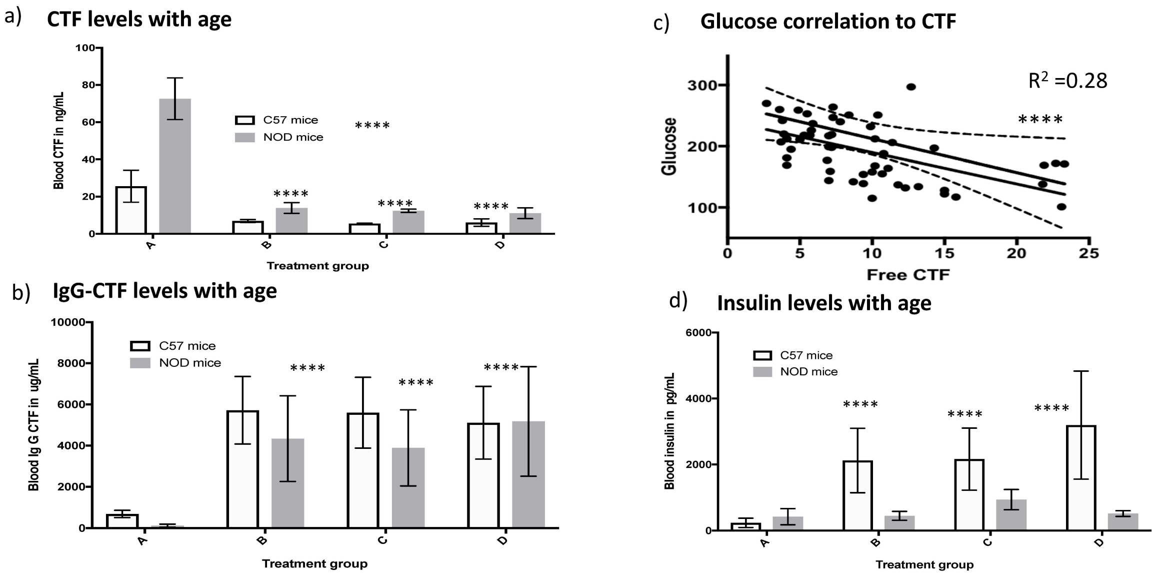


**Supplemental Figure 4. Study 1 blood values shown for Free CTF, Glucose, IgG –CTF and Insulin** for non-obese diabetic mice (NOD) and control mice (C57) on high fat diet. Mice were measured at A) 4-6 weeks of age prior to treatment, B) 16 weeks of age without treatment; C) 16 weeks of age with 10 week of synthetic TACE inhibitor treatment ; D) 16 weeks of age with 10 week of synthetic CTF treatment. **** indicates a P<0.0001.

Glucose tolerance tests in C57 mice showed no significant differences between animals treated with the synthetic TACE inhibitor and those treated with CTF. In contrast, glucose tolerance tests in NOD mice revealed worsening results with 6 weeks of either treatment, suggesting an impact on pancreatic function (Figure S5). Insulin tolerance was not measured in study 1, but this test was added in study 2 to test for the impact of CTF on plasma insulin levels in more detail (**Supplemental Report 2**).

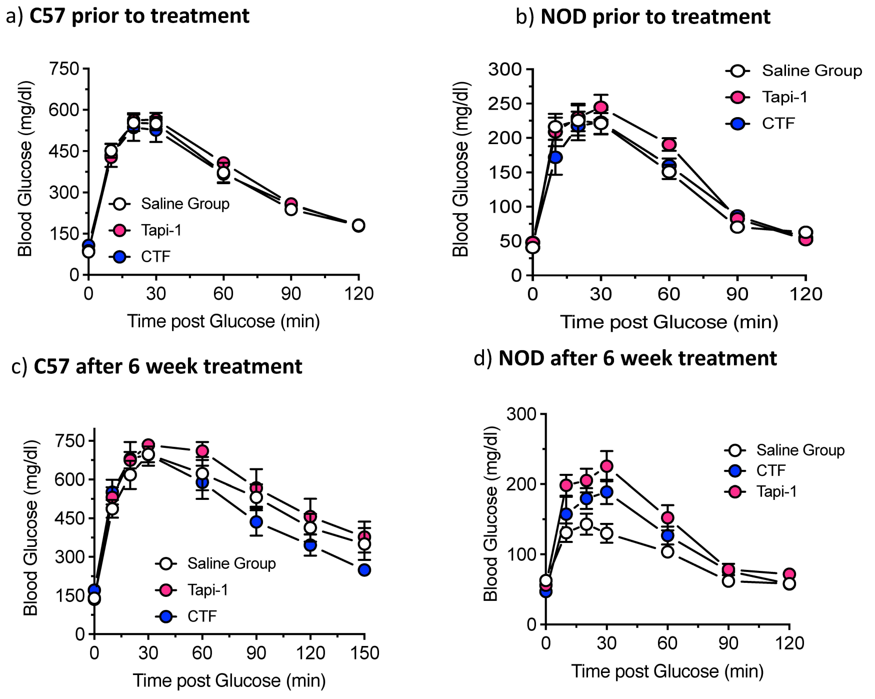


**Supplemental Figure 5. Study 1 Glucose Tolerance Test (ipGTT) data** for control mice (C57) on high fat diet (a & b) and non-obese diabetic mice (NOD) (c & d) at ~6 week of age prior to CTF or TAPI-1 treatment and at ~12 week after weekly treatment. Mice were fasted for either 16 hr (with full access to water) and injected with 2 g/kg glucose by intraperitoneal injection (IP) injection. After dosing with glucose, small (1-2 uL) of blood samples was taken through a tail biopsy at various time intervals (0, 10, 20, 30, 60, 90, 120 and 150 minutes).

The levels of intact adiponectin receptor were measured in the pancreas, liver, brown adipose, muscle and spleen tissue by western blot (Figure S6) and immunocytochemistry (Figure S7) using an antibody specific for the CTF. The most significant changes in the abundance of the intact receptor were observed in the exocrine pancreas followed by the spleen. Western blot analysis revealed that the quantity of intact receptor in the islets of C57 mice was below the level of detection (Figure S6). In NOD mice, the abundance of the receptor declined over time: it was detected in the exocrine pancreas at 6 weeks of age but was undetectable at 16 weeks (Figure S7a). In contrast, the receptor was detected in the exocrine pancreas of C57 mice throughout the experiment, and it accumulated over time (Figure S7c,d). Insulitis (blue) was clearly present around 94% of islets in both untreated and treated NOD mice and in animals treated with TAPI-1, but this declined to only 10% of islets at 16 weeks in NOD mice treated weekly with synthetic exogenous CTF. There was no evidence of insulitis in the C57 mice.

This study suggested that NOD mice provide a good model for adiponectin receptor fragmentation in the pancreas. Given that the treatment of patients with insulitis has merit, we carried out an additional confirmatory study (**Supplemental Report 2**). However, the use of TACE inhibitors was not continued because this did not influence the formation of free CTF.

**Supplemental Figure 6:** Adiponectin receptor expression in C57BL/6 pancreas, spleen, and islet tissue lysate

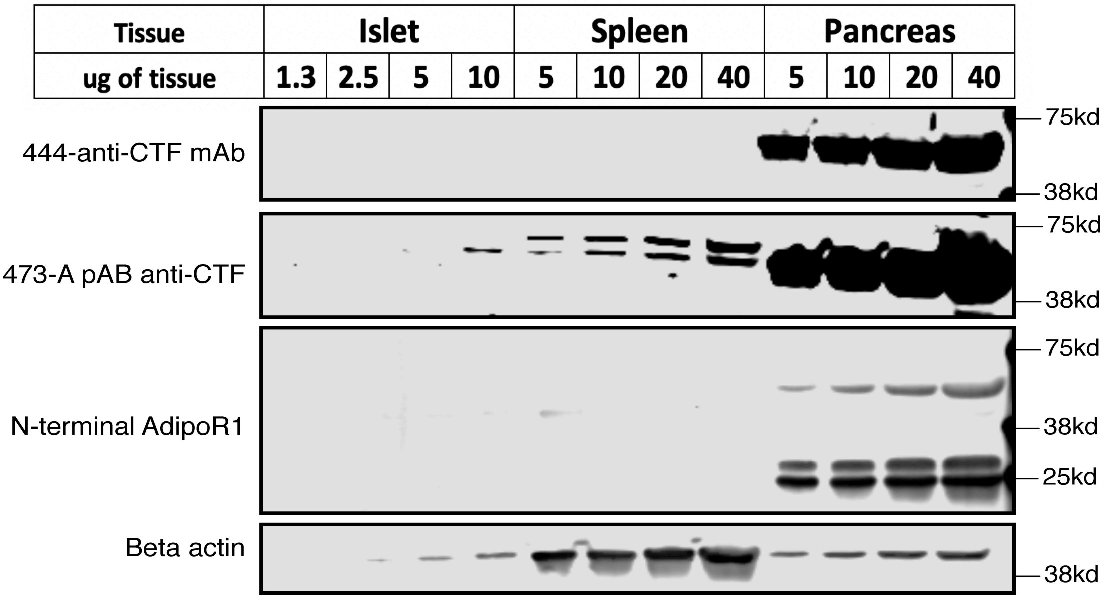


a) **NOD mice on normal diet aged to 16 wk age**

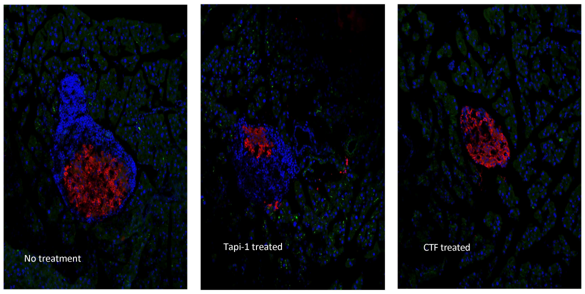


b) **C57 mice on normal diet aged to > 30 wk**

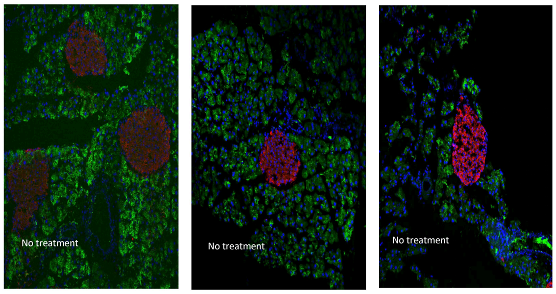


c) **C57 mice on high fat diet aged to 16 wk age**

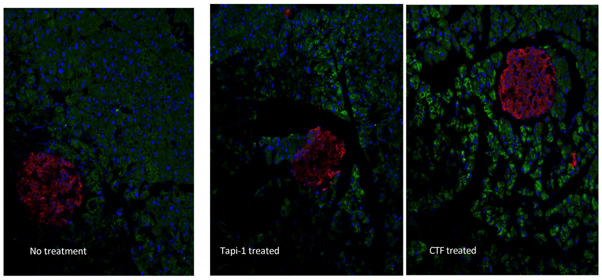


**Supplemental Figure 7. Study 1 ICC Measurements of Pancreatic levels AdipoR CTF and Insulin** for a) non-obese diabetic mice (NOD), b) control mice (C57) on normal or c) high fat diet. Green, AdipoR; Red, insulin; Blue, DAPI Mice were measured at without treatment; with 10 week of synthetic TACE inhibitor treatment or with 10 week of synthetic CTF treatment.

#### Conclusion

The cause of adiponectin receptor fragmentation was not identified in this study, but pancreatic tissue with higher TACE activity caused AdipoR1 fragmentation. The exocrine pancreas also shows pronounced neutrophil and macrophage infiltration as the autoimmune attack on β-cells develops, which would be expected to enhance the proteasome.[6] Bacteria eliciting this adverse innate immunity response in T1D models also add to the proteasome.[7] In agreement, neutrophilic elastase increases in the islets of NOD mice with significant damage (Figure S8).

**Supplemental Figure 8:** Neutrophil Elastase and AdipoR in NOD Islets

Neutrophil Elastase appears to be more prevalent in the hyperglycaemic (terminal, right) mice compared to non-glycaemic mice (Day 105, left)

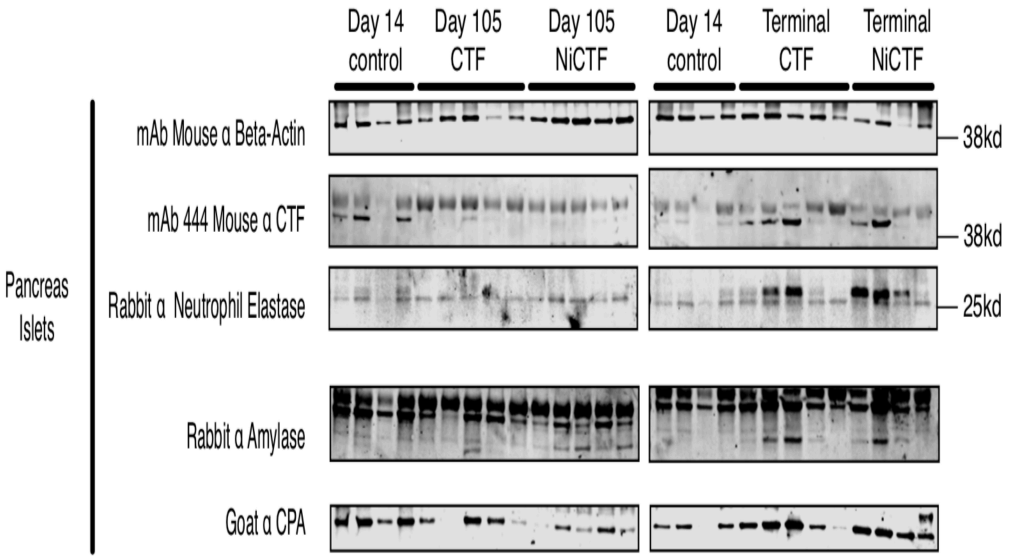


**Acknowledgements**. The Indiana Bioscience Research Institute (an independent non-profit research organization) sponsored the laboratory work behind this study.
