## Supplement Report 2. Antibody blocking of AdipoR CTF inhibition of IDE for "Adiponectin Receptor Fragmentation in Mouse Models of Type 1 and Type 2 Diabetes"

### Blocking the inhibition of IDE by the C-terminal fragment of the adiponectin receptor using a specific autoantibody

#### Introduction

The adiponectin receptor C-terminal fragment (AdipoR1-$CTF_{344\text{-}375}$) is a freely circulating peptide found in the plasma of normal individuals but not in some undefined diabetes patients, a discovery made using a combination of affinity capture and mass spectrometry based on monoclonal antibodies specific for the $CTF_{344\text{-}375}$ peptide.[1,2] The AdipoR1-$CTF_{351\text{-}362}$ peptide domain was identified as a strong non-competitive inhibitor of insulin-degrading enzyme (IDE), whereas the clinical 32-amino-acid AdipoR1 $CTF_{344\text{-}376}$ fragment is a competitive inhibitor of ADAM17 (TACE).[2] Affinity capture proteomics allows the rapid translation of experimental results into novel immunoassays. The concentration of endogenous free AdipoR1-CTF in human peripheral blood is less than <5 ng/mL, which is below the detection limit of sandwich ELISAs, but it can be measured in rodent models.[2] Circulating autoantibodies specific for the AdipoR1-CTF can be detected by sandwich ELISA and its concentration ranges from 5 to 4900 ng/mL in humans and rodents.[38] In normal Sprague Dawley rats, the administration of exogenous AdipoR1-$CTF_{351\text{-}370}$ correlated with increased plasma insulin but this was not the case in Zucker diabetic fatty (ZDF) rats with insulin insufficiency. The interaction between AdipoR1-$CTF_{351\text{-}362}$ and IDE may offer a new therapeutic target because mechanistic and drug studies with IDE have demonstrated an impact on the insulin response.[3-6] We therefore set out to determine whether antibodies specific for AdipoR1-$CTF_{351\text{-}362}$ can block the non-competitive inhibition of IDE by CTF.

#### Experimental (IDE activity inhibition assay)

New lots of the synthetic peptide AdipoR1-$CTF_{351\text{-}375}$ were obtained from Celtek Peptides. The inhibition of IDE activity was determined using the SensoLyte 520 IDE Activity Assay Kit (AnaSpec). IDE activity was determined in the presence of the IDE FRET substrate (SensoLyte ® 520 β - Secretase Assay Kit *Fluorimetric AnaSpec) at concentrations of 1 and 5 μM, 0.1

mg/mL IDE, and increasing concentrations of CTF (0, 0.1, 1 and 10 μM). The reaction kinetics were measured for 2 h at 25°C by excitation at 490 nm (emission = 520 nm) and the slopes for each reaction were calculated. The IDE activity without inhibitor was normalized to 100%. CTF at a concentration of 10 μM reduced IDE activity to 5% in agreement to previous results, and this did not change with different concentrations of the IDE substrate confirming the non-competitive inhibition of IDE by CTF.[1] Two CTF-specific antibodies were tested for their ability to block the inhibition of IDE activity by CTF. Mouse monoclonal antibody 444-1D12 was raised against AdipoR1-CTF$_{351-375}$ and rabbit monoclonal antibody SAT-56-1 was raised against AdipoR1-CTF$_{351-362}$. The light chain and heavy chain DNA sequences for both antibodies are shown in Figures S1 and S2. Control reactions were prepared with whole IgG fraction or BSA in place of the specific antibodies.

Figure S1. Sequence of the mouse monoclonal antibody 444-1D12. (a) Kappa light chain. (b) Heavy chain.

(a)

GATGTTTTGATGACCCAAACTCCACTCTCCCTGCCTGTCAGTCTTGGAGATCAAGCCTCCA
TCTCTTGCAGATCTAGTCAGAACATTTTACATAGTACTGGAAACACCTATTTAGAATGGTAC
CTGCAGAAACCCGGCCAGTCTCCAAAGCTCCTGATCTACAAAGTTTCCAACCGATTTTCTG
GGGTCCCAGACAGGTTCAGTGGCAGTGGATCAGGGACAGATTTCACACTCAAGATCAGCA
GAGTGGAGGCTGAGGATCTGGGAGTTTATTACTGCTTTCAAGGTTCACATGTTCCGCTCAC
GTTCGGTGCTGGGACCAAGCTGGAGCTGAAACGGGCTGATGCTGCACCAACTGTATCCAT
CTTCCCACCATCCAGTGAGCAGTTAACATCTGGAGGTGCCTCAGTCGTGTGCTTCTTGAAC
AACTTCTACCCCAAAGACATCAATGTCAAGTGGAAGATTGATGGCAGTGAACGACAAAATG
GCGTCCTGAACAGTTGGACTGATCAGGACAGCAAAGACAGCACCTACAGCATGAGCAGCA
CCCTCACGTTGACCAAGGACGAGTATGAACGACATAACAGCTATACCTGTGAGGCCACTCA
CAAGACATCAACTTCACCCATTGTCAAGAGCTTCAACAGGAATGAGTGT

(b)

CAGGTTACTCTGAAAGAGTCTGGCCCTGGGATATTGCAGCCCTCCCAGACCCTCAGTCTG
ACTTGTTCTTTGTCTGGGTTTTCACTGAGAACTTCTGGTATGGGTGTGAGCTGGATTCGTCA
GCCTTCAGGAAAGGGTCTGGAGTGGCTGGCACACATTTACTGGGATGATGACAAGAGATA
TAACCCATCCCTGAAGAGCCGGCTCACAATCTCCAAGGATACCTCCAGTAACCAGGTATTC
CTCAAGATCACCAGTGTGGACACTGCAGATACTGCCACATACTACTGTGCTCGAAGACAAA

GCTTTGGTGGCCCCGCTTCTTACGACTGGGGCCAAGGGACTCTGGTCACTGTCTCTGCAG
CCAAAACGACACCCCCATCTGTCTATCCACTGGCCCCTGGATCTGCTGCCCAAACTAACTC
CATGGTGACCCTGGGATGCCTGGTCAAGGGCTATTTCCCTGAGCCAGTGACAGTGACCTG
GAACTCTGGATCCCTGTCCAGCGGTGTGCACACCTTCCCAGCTGTCCTGCAGTCTGACCT
CTACACTCTGAGCAGCTCAGTGACTGTCCCCTCCAGCACCTGGCCCAGCGAGACCGTCAC
CTGCAACGTTGCCCACCCGGCCAGCAGCACCAAGGTGGACAAGAAAATTGTGCCCAGGG
ATTGTGGTTGTAAGCCTTGCATATGTACAGTCCCAGAAGTATCATCTGTCTTCATCTTCCCC
CCAAAGCCCAAGGATGTGCTCACCATTACTCTGACTCCTAAGGTCACGTGTGTTGTGGTAG
ACATCAGCAAGGATGATCCCGAGGTCCAGTTCAGCTGGTTTGTAGATGATGTGGAGGTGC
ACACAGCTCAGACGCAACCCCGGGAGGAGCAGTTCAACAGCACTTTCCGCTCAGTCAGTG
AACTTCCCATCATGCACCAGGACTGGCTCAATGGCAAGGAGTTCAAATGCAGGGTCAACA
GTGCAGCTTTCCCTGCCCCCATCGAGAAAACCATCTCCAAAACCAAAGGCAGACCGAAGG
CTCCACAGGTGTACACCATTCCACCTCCCAAGGAGCAGATGGTCAAGGATAAAGTCAGTCT
GACCTGCATGATAACAGACTTCTTCCCTGAAGACATTACTGTGGAGTGGCAGTGGAATGGG
CAGCCAGCGGAGAACTACAAGAACACTCAGCCCATCATGGACACAGATGGCTCTTACTTC
GTCTACAGCAAGCTCAATGTGCAGAAGAGCAACTGGGAGGCAGGAAATACTTTCACCTGC
TCTGTGTTACATGAGGGCCTGCACAACCACCATACTGAGAAGAGCCTCTCCCACTCTCCTG
GTAAA

Figure S2. Sequence of the rabbit monoclonal antibody SAT-56-1. (a) Kappa light chain. (b) Heavy chain. The IgG variable regions are marked in bold. The DNA fragments feature a HindIII site at the 5' end and a NotI site at the 3' end (underlined italic).

(a)

*AAGCTT*GTACCCTTCACC**ATGGACACGAGGGCCCCCACTCAGCTGCTGGGGCTCCTACT**
**GCTCTGGCTCCCAGGTGCCAGATGTGCTGACATTGTGATGACCCAGACTCCAGCCTCCG**
**TGGAGGCAGCTGTGGGAGGCACAGTCACCCTCAACTGCCAGGCCAGTCAGACCATTGA**
**CGACTACTTATCCTGGTATCAGCAGAAGCCAGGGCAGCCTCCCAAACAACTGATCTACA**
**GGGCATCCACTCTGTCATCTGGGGTCCCATCGCGATTCAAAGGCAGTGGATCTGGGACA**
**GAATTCACTCTCACCATCAGCGCCCTGGAGTGTGCCGATGCTGCCACTTACTACTGTCAA**
**AGTGGTGATTTTAGTGGTGGTCGTAGTTATGGTAATATTTTCGGCGGAGGGACCGAGGTG**
**GTGGTCAAAGGTGAT**CCAGTTGCACCTACTGTCCTCATCTTCCCACCAGCTGCTGATCAG

GTGGCAACTGGAACAGTCACCATCGTGTGTGTGGCGAATAAATACTTTCCCGATGTCACCG
TCACCTGGGAGGTGGATGGCACCACCCAAACAACTGGCATCGAGAACAGTAAAACACCGC
AGAATTCTGCAGATTGTACCTACAACCTCAGCAGCACTCTGACACTGACCAGCACACAGTA
CAACAGCCACAAAGAGTACACCTGCAAGGTGACCCAGGGCACGACCTCAGTCGTCCAGAG
CTTCAATAGGGGTGACTGTTAG

(b)

*AAGCTT*GTACCCTTCACC**ATGGAGACTGGGCTGCGCTGGCTTCTCCTGGTCGCTGTGCTC
AAAGGTGTCCAGTGTCAGTCGCTGGAGGAGTCCGGGGGTCGCCTGGTAACGCCTGGAG
GATTCCTGACACTCACCTGTACAGTCTCTGGAGTCGACCTCAGTGCCTACTGGATGAACT
GGGTCCGCCAGGCTCGTGGGAAGGGGCTGGAGTGGATCGGCACCATTAATCACCGTGG
TAGCACATGGTACCCGAGCTGGGCGAGAGGCCGATTCACCATCTCCAAGACCTCGACCA
CGGTGGATCTGACAATGACCAGTCTGACAACCGAGGACACGGCCACCTATTTCTGTAGT
GGGGGTCTCTTGTGGGGCCCAGGCACCCTGGTCACCGTCTCCTCAGGGCAA**CCTAAGGC
TCCATCAGTCTTCCCACTGGCCCCCTGCTGCGGGGACACACCCAGCTCCACGGTGACCCT
GGGCTGCCTGGTCAAAGGGTACCTCCCGGAGCCAGTGACCGTGACCTGGAACTCGGGCA
CCCTCACCAATGGGGTACGCACCTTCCCGTCCGTCCGGCAGTCCTCAGGCCTCTACTCGC
TGAGCAGCGTGGTGAGCGTGACCTCAAGCAGCCAGCCCGTCACCTGCAACGTGGCCCAC
CCAGCCACCAACACCAAAGTGGACAAGACCGTTGCGCCCTCGACATGCAGCAAGCCCACG
TGCCCACCCCCTGAACTCCTGGGGGGACCGTCTGTCTTCATCTTCCCCCCAAAACCCAAG
GACACCCTCATGATCTCACGCACCCCCGAGGTCACATGCGTGGTGGTGGACGTGAGCCA
GGATGACCCCGAGGTGCAGTTCACATGGTACATAAACAACGAGCAGGTGCGCACCGCCC
GGCCGCCGCTACGGGAGCAGCAGTTCAACAGCACGATCCGCGTGGTCAGCACCCTCCCC
ATCGCGCACCAGGACTGGCTGAGGGGCAAGGAGTTCAAGTGCAAAGTCCACAACAAGGC
ACTCCCGGCCCCCATCGAGAAAACCATCTCCAAAGCCAGAGGGCAGCCCCTGGAGCCGA
AGGTCTACACCATGGGCCCTCCCCGGGAGGAGCTGAGCAGCAGGTCGGTCAGCCTGACC
TGCATGATCAACGGCTTCTACCCTTCCGACATCTCGGTGGAGTGGGAGAAGAACGGGAAG
GCAGAGGACAACTACAAGACCACGCCGGCCGTGCTGGACAGCGACGGCTCCTACTTCCT
CTACAGCAAGCTCTCAGTGCCCACGAGTGAGTGGCAGCGGGGCGACGTCTTCACCTGCTC
CGTGATGCACGAGGCCTTGCACAACCACTACACGCAGAAGTCCATCTCCCGCTCTCCGGG
TAAATGA

### Results

The CTF and CTF-specific antibodies (or control proteins) were pre-mixed and incubated at room temperature for 30 min before adding the IDE and reaction substrate to initiate the reaction. We tested the IDE reaction in the presence of CTF with or without the antibody to determine whether the latter could ameliorate the inhibition of IDE by CFT by forming an affinity complex. We also assessed the reaction kinetics to determine the effect of other reaction components, including the control proteins. We found that a 10-fold molar excess of the CTF-specific antibody almost completely blocked the inhibitory effect of CTF. Other proteins or antibodies present in the mix had no effect on the inhibition of IDE by CTF.

### Conclusion

This study supports the neutralization of plasma IDE inhibition by CTF autoantibodies. We found that antibodies specific for AdipoR1-$CTF_{351\text{-}362}$ were able to block the non-competitive inhibition of IDE by CTF.


### Acknowledgements

The Indiana Bioscience Research Institute (an independent non-profit research organization) sponsored the laboratory work behind this study.
