## Supplement Report 3. Macrophage study of AdipoR CTF Formation for "Adiponectin Receptor Fragmentation in Mouse Models of Type 1 and Type 2 Diabetes"

#### Supplemental Report 3

### The influence of macrophage differentiation and polarization on the formation of the adiponectin receptor C-terminal fragment

Indiana Biosciences Research Institute, Indianapolis IN

##### Introduction

The adiponectin receptor contains an extracellular C-terminal sequence that is released by proteolysis to yield a C-terminal fragment (AdipoR-CTF). The uptake of exogenous CTF by cancer cells *in vitro* was previously shown to increase the levels of intracellular insulin.[1] We set out to investigate whether viable human blood cells in culture would replicate this behavior and whether the activity of tumor necrosis factor-α cleavage enzyme (TACE) could explain the endogenous fragmentation of the receptor. We selected monocytes/macrophages as a model because these cells account for a substantial proportion of the adiponectin receptor found in the blood and TACE is induced during the differentiation of monocytes into macrophages.

##### Adiponectin receptor macrophage signaling

Adiponectin receptor signaling is thought to modulate the behavior of macrophages, dendritic cells and natural killer cells, potentially influencing the process of antigen presentation during the adaptive immune response.[2-4] The downstream signaling pathway involves the phosphorylation of mitogen-activated protein kinase (MAPK p38) to induce pro-hypertrophic signaling in macrophages.[5] A variety of cellular processes are regulated by MAPK p38, including differentiation, apoptosis and responses to inflammation involving the release of cytokines and adhesion molecules.[6] MAPK is activated when the high-molecular-weight form of adiponectin interacts with growth factors, and MAPK signaling inhibits atherogenesis, angiogenesis and proliferation.[7,8] Both the pro-inflammatory behavior of M1 macrophages and the anti-inflammatory behavior of M2 macrophages have been attributed to the adiponectin receptor.[8,9] Adiponectin induces endothelial apoptosis, but also inhibits monocyte adhesion and increasing monocyte lipid uptake during macrophage transformation to foam cells by inhibiting differentiation.[9,7] Adiponectin inhibits the formation of tumor necrosis factor α (TNFα), adhesion molecules, and nuclear factor kappa-light-chain-enhancer of activated B cells (NF-κB) in endothelial cells and macrophages.[10] However, the role of peripheral blood AdipoR in obesity and diabetes remains largely unknown.

**Experimental Design for Macrophage Cultivation**

The differentiation of THP-1 human monocytes into M0 macrophages ($CD63^{+}CD14^{+}$) and their polarization to M1 ($CD206^{+}$) and M2 (insulin receptor $CD220^{+}$) was achieved using phorbol 12-myristate-13-acetate (PMA) with *Escherichia coli* lipopolysaccharide (LPS), interferon gamma (IFNγ), interleukin (IL)-13 and IL-4 as previously reported.[11] The cells were typed by FACS immunoassay on a FACSAria Fusion flow cytometer (BD Biosciences) using nine channels at excitation wavelengths of 405 nm (filters 450/50, 525/50, 610/20), 488 nm (filters 530/30, 659/40), 561 nm (filters 582/15, 610/20, 670/14, 780/60) and 640 nm (filters 670/30, 780/60780/60). The cells were threshold-gated using CD14, CD63, CD220, CD206, AdipoR, IDE, TACE and insulin as previously described.[12,13]

Monoclonal antibodies (mAbs) specific for CD14, CD63, CD63b, CD206, CD220, TACE, IDE and insulin were acquired from commercial suppliers for FACS analysis. Antibodies specific for human CD14 conjugated to Alexa Fluor 647 (mouse mAb, clone #M5E2, 1 μg/μL) and human CD66b conjugated DyLight 650 (mouse mAb, clone #G10F5, 0.1 μg/mL) were purchased from Novus Biologicals. Antibodies specific for human CD63 conjugated to phycoerythrin (PE) (mouse mAb, cat #556020, 0.2 μg/μL), human CD206 conjugated to PE (mouse mAb, clone #19.2, 0.2 μg/μL), human CD220 conjugated to BD Horizon BB700 (mouse mAb, clone #3B6/IR, 0.2 μg/μL), and a mouse IgG1κ isotype control conjugated to PE (mouse mAb, clone #559320, 0.2 μg/μL) were purchased from BD Biosciences. Antibodies specific for human IDE conjugated to Alexa Fluor 647 (mouse mAb, clone #334501, 1 μg/μL), human TACE conjugated to Alexa Fluor 594 (mouse mAb, clone #136133, 1 μg/μL), and mouse IgG conjugated to Alexa Fluor 647 (mouse mAb, clone #11711, 1 μg/μL) were purchased from R&D Systems. Antibodies specific for human insulin conjugated to Alexa Fluor 488 (mouse mAb, clone #ICBTACLS, 0.5 μg/μL) were purchased from Thermo Fisher Scientific. Antibody concentrations were determined by Nanodrop spectrophotometry based on an approximate molecular weight of 150 kDa. The Fc receptor on monocytes and macrophages was blocked with IgG and verified using a mouse IgG1κ isotype control.[14] A mouse mAb specific for AdipoR1-$CTF_{351\text{-}375}$ (clone 444-1D12-1H7) was produced from hybridoma cell lines (ATCC) and conjugated to FITC (IBRI, Indianapolis, IN). Cell lysates were tested for TACE activity using the SensoLyte 520 TACE Activity Assay Kit (AnaSpec). TNFα alpha was used as a proxy for TACE activation, and levels were determined using an Invitrogen Human TNFα Uncoated ELISA kit (Thermo Fisher Scientific, 88-7346-88, Lot# 168722002).

The transformation and differentiation of human monocytes (THP-1) to M0, M1 and M2 macrophages was carried out as follows.[11] THP-1 cultures were seeded from –80°C stock vials ($1\times10^{6}$ cells/mL) into T-25 flasks containing 10 mL RPMI 1640 medium with phenol red (Thermo Fisher Scientific) adjusted with 10% fetal bovine serum, 10 mM HEPES, 1 mM sodium pyruvate,

2.5 g/L D-glucose (Thermo Fisher Scientific), 50 pM β-mercaptoethanol and 1% penicillin-streptomycin (PenStrep, Sigma-Aldrich). The insulin concentration in 10% FBS is 80 pM. The cells were incubated at 37°C in a 5% $CO_2$ atmosphere until they reached a density of $2\times10^6$ cells/mL, typically within 3–4 days.

Differentiation to M0 macrophages was induced by incubating the monocytes in medium containing 200 nM PMA (Sigma-Aldrich, P8139) for 24 h followed by resting for 24 h, by which time larger M0 cells can be visualized by phase contract microscopy. Differentiation to M1 was induced over a period of 124 h by adding 20 ng/mL recombinant human INFγ (R&D Systems, 285-IF-100) and 10 pg/mL LPS (Sigma-Aldrich, L4391). Differentiation to M2 was induced over a period of 24–72 h by adding 20 ng/mL IL-4 and 20 ng/mL IL-13 (Fig 2). Differentiation to M0, M1 and M2 macrophages was verified by threshold gating using antibodies specific for CD14, CD63, CD63b CD206, and CD220.[12,13]

Cells were collected by transferring the medium to Falcon tubes and incubating the cells with 2 mL Accutase (Sigma-Aldrich, A6994) for 5 min at room temperature. The detached cells were then transferred to the Falcon tubes containing the aspirated medium. The concentration of cells was measured, and then the tubes were centrifuged at 200 × *g*, the supernatant was removed, and the cells were resuspended in an appropriate volume of PBS to achieve a concentration of $5\times10^6$ cells/mL. The cells were centrifuged at 200 × *g* for 10 min, washed in 100 µL PBS and centrifuged as above. Cell viability was then determined by resuspending them in 100 µL PBS containing 1 µL of reconstituted LD violet stain (50 µL DMSO/vial; Thermo Fisher Scientific, L34963) and incubating for 30 min at room temperature. We compared trypan blue, propidium iodide, 4′,6-diamidino-2-phenylindole (DAPI) and calcein AM with LD violet, revealing that the latter was most suitable for the selection of live cells over necrotic and apoptotic cells by flow cytometry. The cell viability was >95%.

The cells were washed with 100 µL PBS and incubated for 15 min in 100 µL PBS containing 0.5% BSA (Sigma-Aldrich, 05470), 0.1% Tween-20 (Sigma-Aldrich, P1379) and 10 µg/mL human IgG (Sigma-Aldrich, I4506) to block Fc binding. The blocking agent was then removed and the cells were incubated for 30 min at room temperature in 100 µL PBS containing 1 µL of the antibodies specific for CD14 CD63, CD66b, CD200, TACE, IDE, AdipoR CTF or insulin, labeled with florescent dyes as described above. After a further wash with PBS containing 1% BSA, the cells were fixed with 3.7% paraformaldehyde (PFA) in PBS for 15 min followed by another wash with PBS containing 1% BSA. Finally, the cells were resuspend in 200 µL PBS for FACS analysis. Cell supernatants were prepared by centrifuging cells at 1200 × *g* for 10 min, removing the supernatant and adding protease inhibitor prior to storage at –70°C. Cell lysates were prepared by washing the cells in PBS, and resuspending them in 200 µL of the appropriate lysate buffer followed by incubation on ice for 20 min. The suspension was then centrifuged at 10,000 × g for 10 min at 4°C, and the supernatant was stored at –70°C.

FACS analysis revealed a shift from THP-1 monocytes (CD14$^{high}$ CD63$^{low}$ CD206$^{low}$ CD220$^{high}$) to M0 macrophages (CD14$^{low}$ CD63$^{high}$ CD206$^{low}$ CD220$^{low}$) as shown in Figure S1. CD14 is a membrane receptor for LPS, which is upregulated on macrophages. CD63 (lysosome-associated membrane protein 3) is present in the endosomes and plasma membrane of macrophages and is involved in phagocytosis. CD206 is the macrophage mannose receptor and CD220 is the insulin receptor. M0 macrophages shifted to CD14$^{high}$ CD63$^{low}$ CD206$^{low}$ CD220$^{high}$ when polarized to the M1 phenotype, and to CD14$^{low}$ CD63$^{low}$ CD206$^{high}$ CD220$^{low}$ when polarized to the M2 phenotype.

The effect of the AdipoR1-CTF$_{351-375}$ peptide and the TACE inhibitor TAPI-1 was tested by adding these reagents to the THP-1 monocytes before differentiation to M0 macrophages. A 50 mg/mL solution of AdipoR1-CTF$_{351-375}$ (Celtek Peptides) or TAPI-1 (Sigma-Aldrich, SML0739) was diluted 1:10 in RPMI 1640 medium containing 10% DMSO followed by dilution to 1 mg/mL in medium containing 2% DMSO and filter sterilization. A control with no CTF or TAPI-1 was prepared in medium containing 2% DMSO or 2% DMF.

**Results**

Monocytes and macrophages were analyzed by FACS to determine the numbers of cells producing TACE, IDE, insulin and the adiponectin receptor (Figure S1). The CD63$^{+}$ (M0) macrophages remained viable following differentiation induced by PMA (Figure S1a,b). Following differentiation, we observed a fall in the proportion of cells displaying the intact adiponectin receptor or containing insulin, but an increase in the number of cells containing IDE (Figure S1c,d). Specifically, we found that 30–40% of the monocytes were positive for the receptor but only 10–20% of the M0 macrophages, and that 20–30% of the monocytes contained insulin but this fell to 0–10% in the macrophages. In contrast, 50–55% of monocytes contained IDE but this increased to 80–90% in the macrophages. Overall there were fewer TACE$^{+}$ cells but a greater change during differentiation, rising from 0.5% among the monocytes to 4.5% in the macrophages. Exposing the cells to exogenous AdipoR1 CTF$_{351-370}$ or TAPI-1 had no discernable effect.

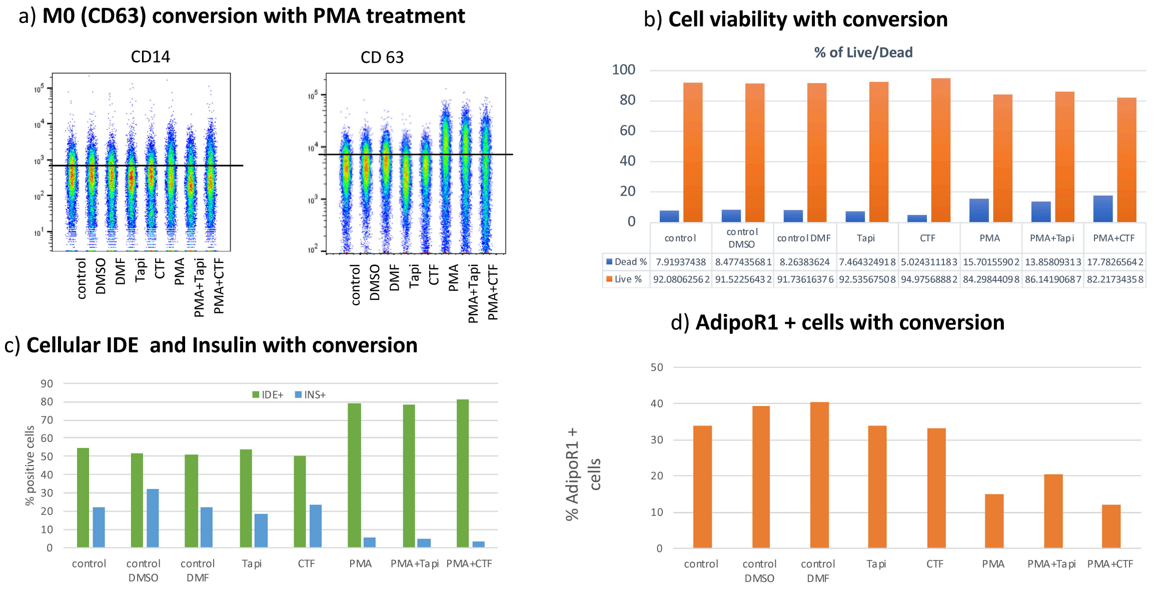


**Supplemental Figure 1. Conversion from THP-1 monocytes to M0 macrophages** are shown for FAC analysis after a) immunocytochemical (ICC) staining for a) CD14 and CD 63 b) DAPI staining for viability c) ICC staining for insulin and IDE positive cells, and d) for AdipoR1 cells with intact CTF.

Western blots confirmed the depletion of the intact adiponectin receptor in M0, M1 and M2 macrophages, with concurrent increases in TACE activity, MAPK phosphorylation and the expression of AMPK (Figure S2). There was little difference between the M0, M1 and M2 macrophage phenotypes (Figure S2). The macrophages remained viable after exposure to AdipoR1 $CTF_{351-370}$ or the TACE inhibitor TAPI-1, and there was no effect on the proportions of cells positive for IDE, the adiponectin receptor, or insulin. Additionally, exogenous AdipoR1-$CTF_{351-375}$ was not taken up by the viable cells, as determined by a sandwich ELISA for the detection of free CTF in cell lysates (LoD = 0.267 ng/mL, LoQ = 0.372 ng/mL).

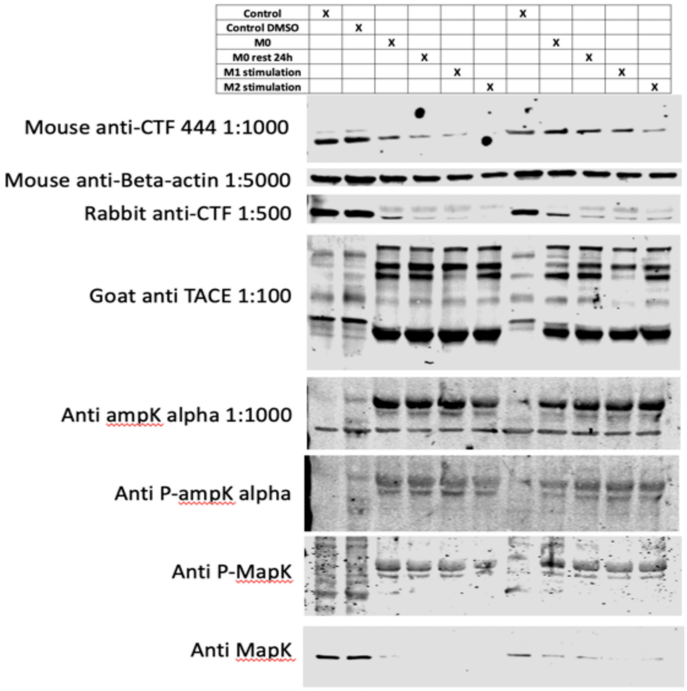


**Supplemental Figure 2.**

Western blot confirmed reduced intact AdipoR CTF upon activation to M0/M1/M2 stages and also showed increase in TACE, MAPK phosphorylation and increase expression of AMPK

The release of endogenous free CTF was detected by sandwich ELISA following macrophage polarization from M0 to M1 (Figure S3a,b). CTF cleavage occurred during M1 polarization, along with the production of TNFα (Figure S3c) but there was no concomitant increase in TACE activity (Figure S3d). Activity assays require the release of TACE from the membrane to allow substrate cleavage, whereas the TNFα release assay can distinguish between inactive and active membrane-bound TACE.

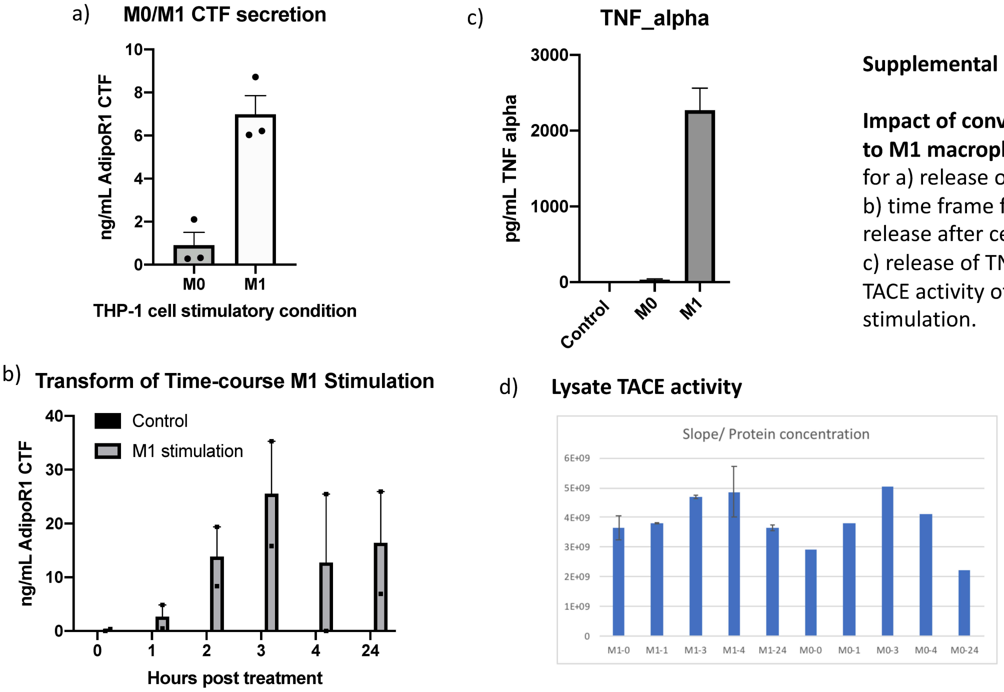


**Supplemental Figure 3.**

**Impact of conversion from M0 to M1 macrophage** are shown for a) release of AdipoR1 CTF; b) time frame for AdipoR1 CTF release after cell stimulation; c) release of TNF- alpha and d) TACE activity of lysates after cell stimulation.

#### Conclusions

Viable human blood cells in culture did not take up exogenous CTF, and no direct impact on intracellular insulin was observed. TACE activity did not explain the endogenous fragmentation of the receptor, but we observed a correlation with TNFα release, and endogenous fragmentation of the receptor occurred following the polarization of M0 to M1 macrophages.

#### Acknowledgements

The Indiana Bioscience Research Institute (an independent non-profit research organization) sponsored the laboratory work behind this study.
